# Overexpression of Active or Inactive RNase J Disrupts Chloroplast RNA Quality Control in Arabidopsis

**DOI:** 10.64898/2026.10.05.756640

**Authors:** Sarah Gartner, Clara Bougearel, Gadi Schuster, Benoît Castandet, David B. Stern, Kevin Baudry

**Author notes:** To whom correspondence should be addressed. (KB), (DBS).

## Abstract

The chloroplast is a semi-autonomous organelle with a compact genome encoding ∼100 expressed genes. Numerous and widespread promoters coupled with inefficient transcription termination leads to a highly complex primary transcriptome that undergoes extensive RNA quality control orchestrated by RNA binding proteins and ribonucleases. Among the ribonucleases, only RNase J (RNJ) is absolutely required for plant germination and survival, likely because its deficiency leads to massive accumulation of antisense RNA which inhibits translation. As one approach to understanding the selectivity of RNJ for antisense RNA and noncoding regions, we created Arabidopsis lines overexpressing catalytically inactive versions of the enzyme designed to trap its substrates. Initially, transgenic plants appeared phenotypically normal. However, crosses used to generate homozygous transgenic lines yielded progeny that developed varying degrees of chlorosis within 2-3 weeks. Concomitantly, these plants exhibited chloroplast transcriptome alterations similar to RNJ-silenced lines, along with mRNA changes consistent with retrograde signaling. This chlorosis could be correlated to the likely decay of RNJ protein, but not its endogenous transcript. The result could be replicated by attempting to overexpress a wild-type version of RNJ, in which case even one transgene copy was sufficient to induce chlorosis. Our results suggest that RNJ is under protein expression level surveillance to prevent wholesale destruction of the plastid transcriptome.

## INTRODUCTION

Land plant chloroplast genomes encode roughly 100 protein and RNA products, whose collective expression is essential for embryogenesis and plant development. While derived from a cyanobacterial endosymbiont, the genome has adapted to its eukaryotic context through transfer of most genes to the nucleus, including those encoding all proteins required for maturation of the transcriptome. These factors act to cleave, trim, splice, modify and edit the myriad of primary transcripts that arise from an imprecise system with hundreds of promoters on both DNA strands, coupled with what appears to be stochastic transcription termination (Small *et al*., 2020; Rovira and Smith, 2019; Castandet *et al*., 2019; Zhelyazkova *et al*., 2012; Wang *et al*., 2022).

There are several classes of ribonucleases (RNases) in the chloroplast, namely endoribonucleases, exoribonucleases and a single protein, RNase J (RNJ), that is a bifunctional endo-exoribonuclease. Dual RNJ functions can both be robust as in *Arabidopsis* (Halpert *et al*., 2019) or unbalanced, with the enzyme found in *B. subtilis* being primarily exonucleolytic (Mathy *et al*., 2007), and several other RNJs from single-celled organisms being endonucleolytic (Levy *et al*., 2011; Liponska *et al*., 2018). RNJ is also unique in that unlike other chloroplast RNases, it is required for embryo development in *Arabidopsis* (Chen *et al*., 2015) and possibly in other plant species. This suggests that unlike other chloroplast RNases, it is both non-redundant and required for sufficient expression of at least one of the chloroplast genes required at this developmental stage.

Due to embryo lethality of *rnj* null mutants, functional dissection of plant RNJ has relied on *in vitro* analysis of recombinant proteins combined with RNA analysis of plants where RNJ expression has been reduced by virus-induced gene silencing (VIGS). The former approach identified the two nucleolytic activities as mentioned above, and investigated a plant-specific C-terminal domain related to GT-1 transcription factors, that was subsequently found to assist in dsRNA binding and localization of RNJ to the plastid nucleoid (Baudry *et al*., 2026). VIGS had previously shown in both tobacco and *Arabidopsis* that RNJ deficiency leads to massive accumulation of antisense RNA (asRNA) that can interfere with sense strand translation, as well as incomplete maturation of several mRNA 5’ termini that depend on RNJ’s exonucleolytic function (Sharwood *et al*., 2011; Luro *et al*., 2013). Since a number of chloroplast asRNAs normally accumulate in wild-type (WT) plants (Hotto *et al*., 2010; Hotto *et al*., 2011; Zghidi-Abouzid *et al*., 2011; Georg *et al*., 2010), this raised the question of whether and if so how, RNJ exerts selectivity over the primary transcript population.

One approach to testing for selectivity would be co-immunoprecipitating a catalytically disabled RNJ with its substrates. The concept of using cleavage-defective RNases to do so has been demonstrated previously for bacterial RNase III (Lioliou *et al*., 2012; Gatewood *et al*., 2012) and human RNase H1 (Chen *et al*., 2017). RNase J’s catalytic mechanism has been well explored using structural approaches (Dorléans *et al*., 2011; Zhao *et al*., 2015; Bao *et al*., 2023) and among its conserved segments is the metallo-β-lactamase (MBL) domain within which lie seven highly conserved motifs, including motif II with the consensus sequence HxHxDH. These residues are involved in coordinating zinc ions that participate in catalysis (Pei *et al*., 2015) and mutation of either the terminal Asp or His in the *Deinococcus radiodurans* enzyme largely abolished *in vitro* catalytic activity (Zhao *et al*., 2015), whereas mutating these two residues along with the preceding His had the same effect on the *Arabidopsis* enzyme (Halpert *et al*., 2019). Thus, motif II is an appropriate target for expressing a catalytically inactive version of RNJ *in vivo*.

We have recently shown that it is possible to complement the *Arabidopsis rnj* null mutant with versions of RNJ lacking the C-terminal GT1 domain (Baudry *et al*., 2026). This is possible because GT1 mutations do not fully abrogate enzyme activity and thus the plants are able to complete a developmental cycle. Since motif II mutations are expected to inactivate the enzyme and thus cause embryo lethality, it was likely that co-expression with a WT version would be required to support pull-down approaches. Below, we show that motif II mutant proteins can indeed be expressed only in the presence of WT RNJ, but that transgene copy number determines whether plants survive or undergo a progressive chlorosis correlated with likely post-transcriptional silencing of the endogenous *RNJ* gene. We also found that when WT RNJ was expressed from a strong 35S promoter chlorosis and silencing also resulted, whereas transgenic expression from the native *RNJ* promoter allowed complementation of the mutant. These results suggest that the expression level of RNJ itself is critical for chloroplast function, and that excess RNJ triggers a retrograde signal that targets the nuclear *RNJ* locus.

## RESULTS

### Transgenic lines expressing catalytically inactive RNJ segregate chlorotic progeny

Our initial goal was to identify RNA targets of RNJ by immunoprecipitating RNJ and sequencing bound RNAs, using catalytically inactive RNJ to avoid cleaving captured RNAs. We created two inactive versions of RNJ, targeting motif II from within the MBL domain, as discussed above and detailed in Figure S1. To ensure that the mutations would inactivate RNJ in our system, we copied the triple mutation used for *in vitro* studies of *Arabidopsis* RNJ (Halpert *et al*., 2019), replacing three residues in the conserved motif HxHxDH with Ala or Lys (Figure S1b). These mutations were placed into two contexts: full-length RNJ, and RNJ lacking the C-terminal GT1 domain (Figure 1a). Our logic was that by comparing results with the two constructs, RNAs dependent on the dsRNA binding activity of the GT1 domain could be distinguished from those binding to the catalytic site.

**Figure 1.**
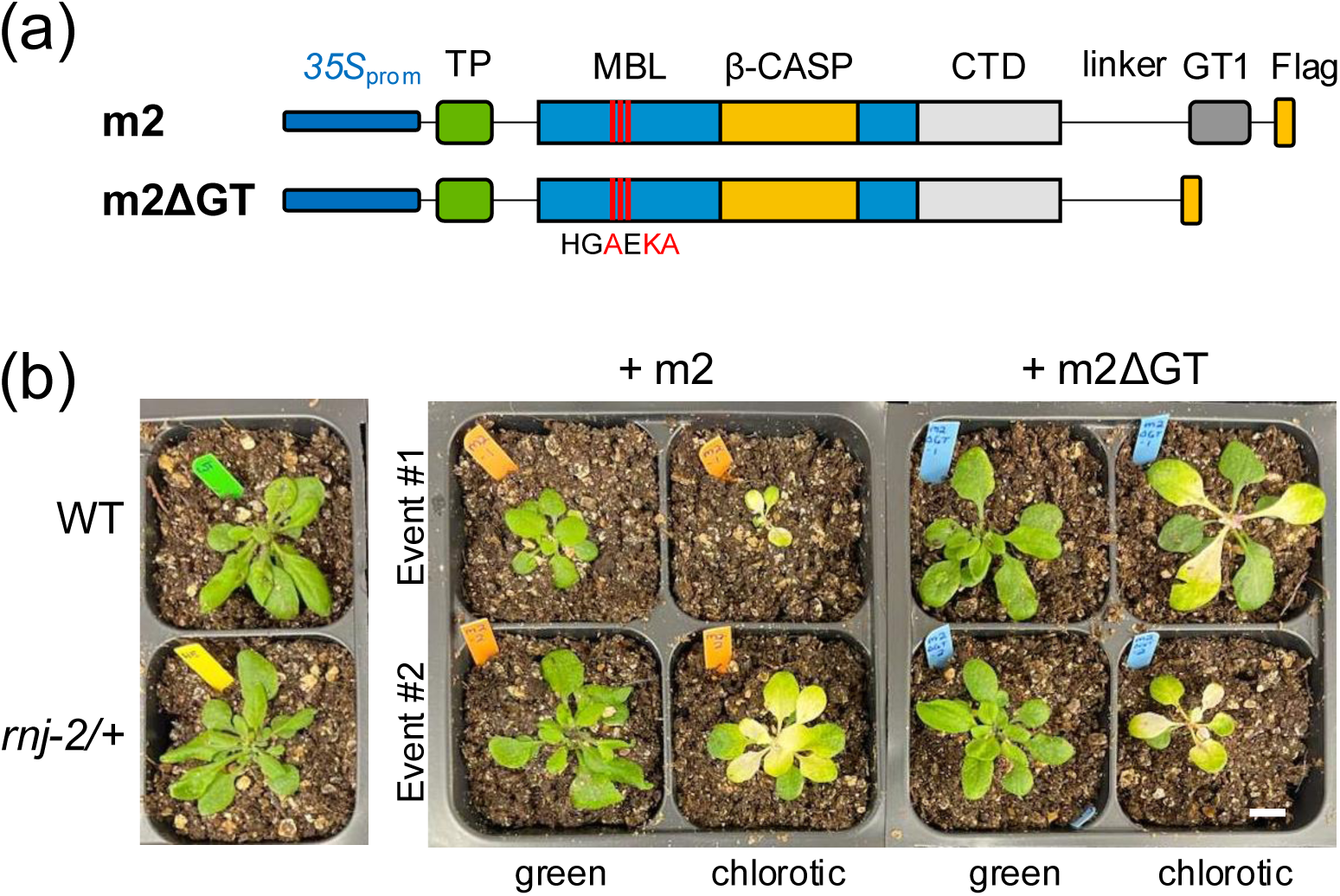
Transgenes and growth phenotypes of *motif II* lines. (a) Constructs used to transform the heterozygous *rnj-2* mutant. The CaMV 35S promoter was used to drive expression of motif II mutant RNJ with or without a GT1 region, flanked by a Flag epitope tag. Domains drawn to scale are the TP, chloroplast target peptide; MBL, Metallo-β-lactamase domain; β-CASP, β-CASP domain; CTD, C-terminal domain; GT1, GT1 domain. Red lines indicate mutated residues of motif II as indicated in the sequence below. (b) Top left, untransformed WT Arabidopsis; bottom left, heterozygous *rnj-2/+* mutant. Right side of panel, representatives of green and chlorotic populations from two independent transformation events (Event 1, top row; Event 2, bottom row) of the indicated constructs. Plants were grown for one month in long day conditions at 22°C (scale bar = 1 cm).

Our previous studies of the GT1 domain used constructs driven by the native *RNJ* promoter for complementation of the *rnj-2* T-DNA insertional mutant (Baudry *et al*., 2026). For the current study we instead chose the strong 35S promoter, reasoning that higher expression of RNJ might provide more robust recovery of RNA. As before, the coding region was flanked by Flag tag to facilitate identification in plant extracts. These constructs will be referred to respectively as m2 (the full-length version) and m2ΔGT (the GT1 truncated version). We transformed heterozygous *rnj-2/+* plants and screened the T1 generation for presence of the transgene, which would be hemizygous in this generation. We also used PCR to screen for the *rnj-2* T-DNA insertion, seeking heterozygotes that upon selfing would generate the three possible genotypes at that locus. For each construct, two independent insertion events were propagated.

T1 plants were virtually identical to WT or *rnj-2/+* plants, that is, they appeared to grow normally and to maturity. In their progeny (the T2 generation), however, two plant populations were observed in all four cases (two events x two constructs): one green and one chlorotic, as exemplified in Figure 1b. Among these T2 plants, none was identified as homozygous for the *rnj-2* T-DNA insertion, which was expected given that expression of inactive RNJ should not complement the known embryo lethality of the mutant background (Chen *et al*., 2015). In addition, both chlorotic and green plants were found to be either wild-type *RNJ/RNJ* or heterozygous *rnj-2/+*, meaning that the phenotype is not dependent on the presence of the *rnj-2* T-DNA insertion. Because the phenotype was observed for four independently selected lines, position effects of transgene insertion site can be ruled out. From this point, we focused on the highly unexpected chlorosis phenomenon rather than recovery of bound RNA, which will be pursued in a future study.

### Chlorosis arises during plant development and is non-Mendelian

To characterize the chlorotic plant populations more systematically, we observed seedling phenotypes for 19 days after sowing (DAS). We noted that upon germination on plates, all seedlings were green and at a gross level indistinguishable from WT plants. After the emergence of the first true leaves at roughly 13-14 DAS, some seedlings began to develop chlorosis at the center of their rosette, that eventually spread across the same leaf and propagated to the subsequent emerging leaves (Figure 2a). Over time, chlorotic areas began to bleach, and as is typical in variegation situations where plants are maintained in the light, was irreversible (Yu *et al*., 2007). Most plants that failed to develop chlorosis by 14 DAS remained green and indistinguishable from WT plants (see below).

**Figure 2.**
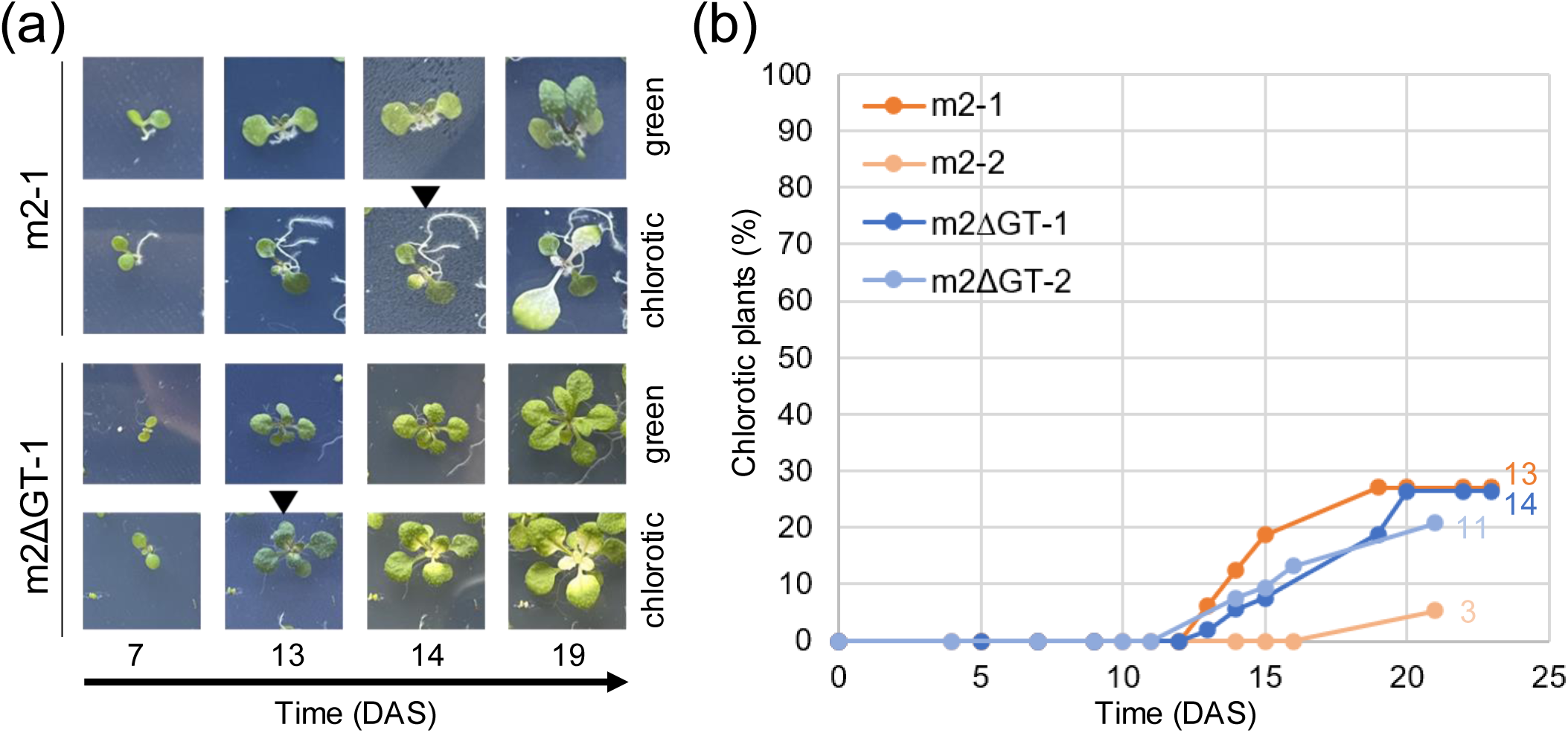
Chlorosis development in *motif II* lines. (a) Time course of representative m2-1 (top two rows) or m2ΔGT-1 (bottom two rows) plants that either remain green or develop chlorosis. For each series, the same plant is followed over time, with days after sowing (DAS) is indicated at the bottom of each column. Black arrowheads indicate the first day that chlorosis was noted. (b) Percentage of all plants that developed chlorosis over the 19-day time course. Numbers at the end of each curve indicate the total number of chlorotic plants observed.

Plants developing chlorosis continued to grow, albeit more slowly than green plants. In the case of m2ΔGT lines, most chlorotic plants were able to achieve a full life cycle and yield viable seeds. In contrast, chlorotic plants of the m2 lineage rarely bolted and thus did not produce seeds. To estimate the proportion and the timeline of chlorosis emergence, we sowed about 90 seeds of each event on media and followed their individual development for several weeks. Most chlorosis initiated within about 15 DAS, with occasional emergence up to 21 DAS. At the end point of our observations, approximately 25% of plants were chlorotic for three of the four events, with line m2-2 being an outlier and consistently generating few chlorotic plants (Figure 2b).

### Chlorotic phenotype correlates with transgene copy number

While the *rnj* genotype could not explain the development of chlorosis, ploidy of the transgene was another possibility. The T2 population, when germinated on hygromycin to select for at least one transgene, in principle harbors 1/3 transgene homozygotes and 2/3 hemizygotes. A 1/3 ratio is close to the ∼25% chlorosis observed for most lines. The copy number of the transgene was evaluated using qPCR in green and chlorotic plants in the four lines, with primers specific to the 3’ UTRs of the native *RNJ* gene and transgene. Because the *rnj-2* T-DNA is in the second intron, it is far upstream of the amplicon and thus plants with *RNJ/RNJ* or *rnj/RNJ* genotypes will behave the same (Figure S2a).

Figure 3 shows that compared to green plants, chlorotic ones have twice as many transgene copies, ranging from 1.9 to 2.2 times. This figure was derived by dividing the signal for the transgene by that of the native gene, given that the endogenous *RNJ* gene is single copy. Consequently, the ratio of 0.5 for the green plants indicates a single transgene in the hemizygous state. The ratio of 1 for the chlorotic plants could result either from a single homozygous insertion, or from two hemizygous insertions. Since we have never observed a copy number result suggestive of more than two copies, we conclude that a single homozygous insertion is the most likely explanation. In line m2ΔGT-2 where the ratio is >1.5, it is possible that there is a complex insertion that results in a higher qPCR signal. In this experiment, we again verified that the results were independent of the *rnj-2* mutant background (Figure S2b). The phenotype and the transgene copy number are therefore correlated, but independent of the *RNJ* genotype.

**Figure 3.**
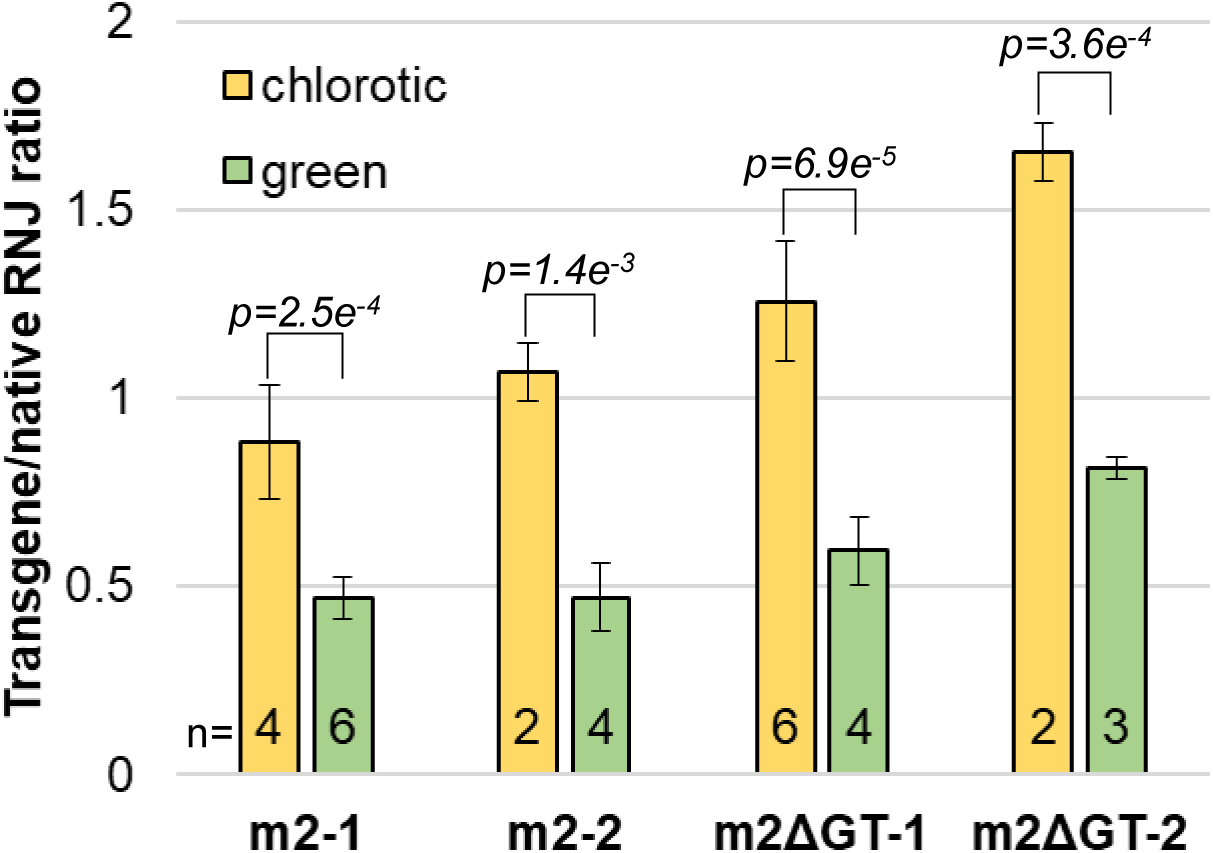
Transgene copy number is associated with the T2 plant phenotype. Evaluation of transgene copy number in chlorotic (yellow bars) and green (green bars) T2 plants by qPCR. Transgene Ct values are normalized to those obtained for the native locus of *RNJ*. *p* values are T-test p-values. The number of plants analyzed for each bin (n) is shown at the bottom of each bar.

To further confirm this result and test its heritability, we characterized progeny of a chlorotic m2ΔGT-1 plant from the T2 generation, in which all plants should be homozygotic transgenics. The progeny (T3 generation) were all green after germination, however 100% of the plants developed chlorosis in the subsequent 2-3 weeks (Figure S3a) and have similar inferred transgene copy numbers to the parent (Figure S3b). Altogether, our results confirm that chlorotic plants, but not persistently green plants, are homozygous for the transgene insertion.

### Chloroplast RNA quality control is impaired in chlorotic plants

Chlorosis can result from disruption of many different processes and/or gene expression changes. Because the phenotype studied here correlated with the presence of RNJ transgenes, we examined whether underlying the chlorosis were molecular signals of RNJ deficiency. VIGS of Arabidopsis and tobacco plants targeting *RNJ* causes chlorosis, with the most striking molecular phenotype being the massive accumulation of asRNA that form sense-antisense duplexes (Sharwood *et al*., 2011). To quickly survey whether the *motif II* lines also accumulate such transcript, we probed total RNA with an anti-dsRNA antibody in a dot blot analysis. For WT, and green m2-1 and m2ΔGT-1 plants no signal was observed, suggesting a low level of dsRNA (Figure 4). In contrast, the strong signal observed for chlorotic plants indicates a much higher level of dsRNA accumulation, presumably in the chloroplast. We are not aware of any other mutants affecting chloroplast RNA metabolism that result in this phenotype.

**Figure 4.**
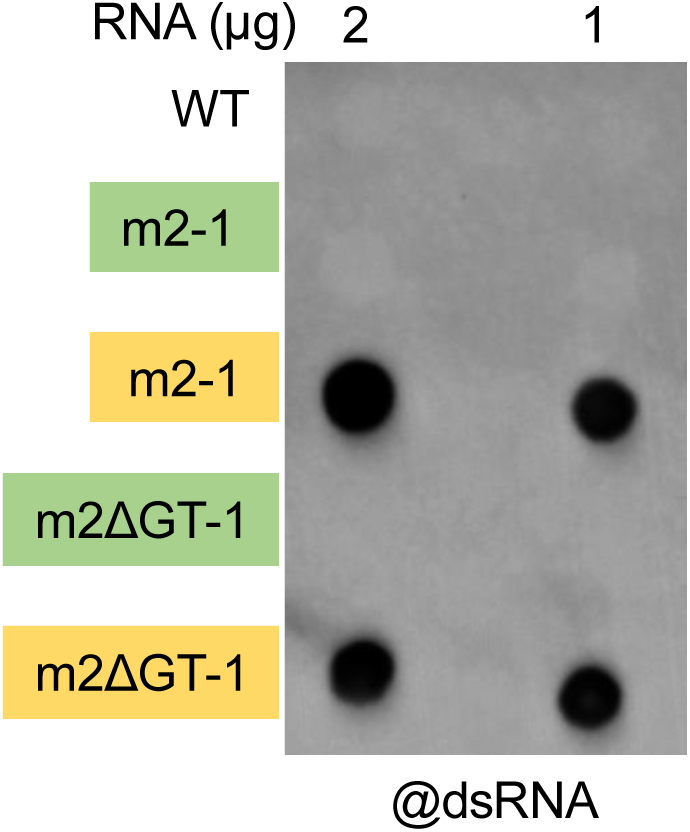
Chlorotic plants accumulate dsRNA. Green and yellow boxes indicate green and chlorotic plants, respectively. Total RNA was extracted from the indicated lines, and 2 or 1 µg was applied to a dot blot, which was probed with the J2 anti-dsRNA antibody.

To assess the state of the plastid transcriptome more directly, we used gel blots and 5’ RACE. Total RNA was isolated and as expected, chlorotic plants accumulated less plastidial rRNA (16S and 23S), whereas RNA from green plants displayed a WT rRNA pattern (Figure 5a). Because chloroplast rRNA quantitative or qualitative changes can occur for many different reasons (Schmid *et al*., 2024), we used gel blot hybridization to test for presence of asRNA to *psbI*, which was one of the hallmarks of RNJ deficiency observed in the VIGS study. As shown in Figure 5b, as-*psbI* is readily detected in RNA from all chlorotic plants, whereas no signal was observed for the WT and green *motif II* plants. This result further suggested that RNJ function was at play in the development of chlorosis.

**Figure 5.**
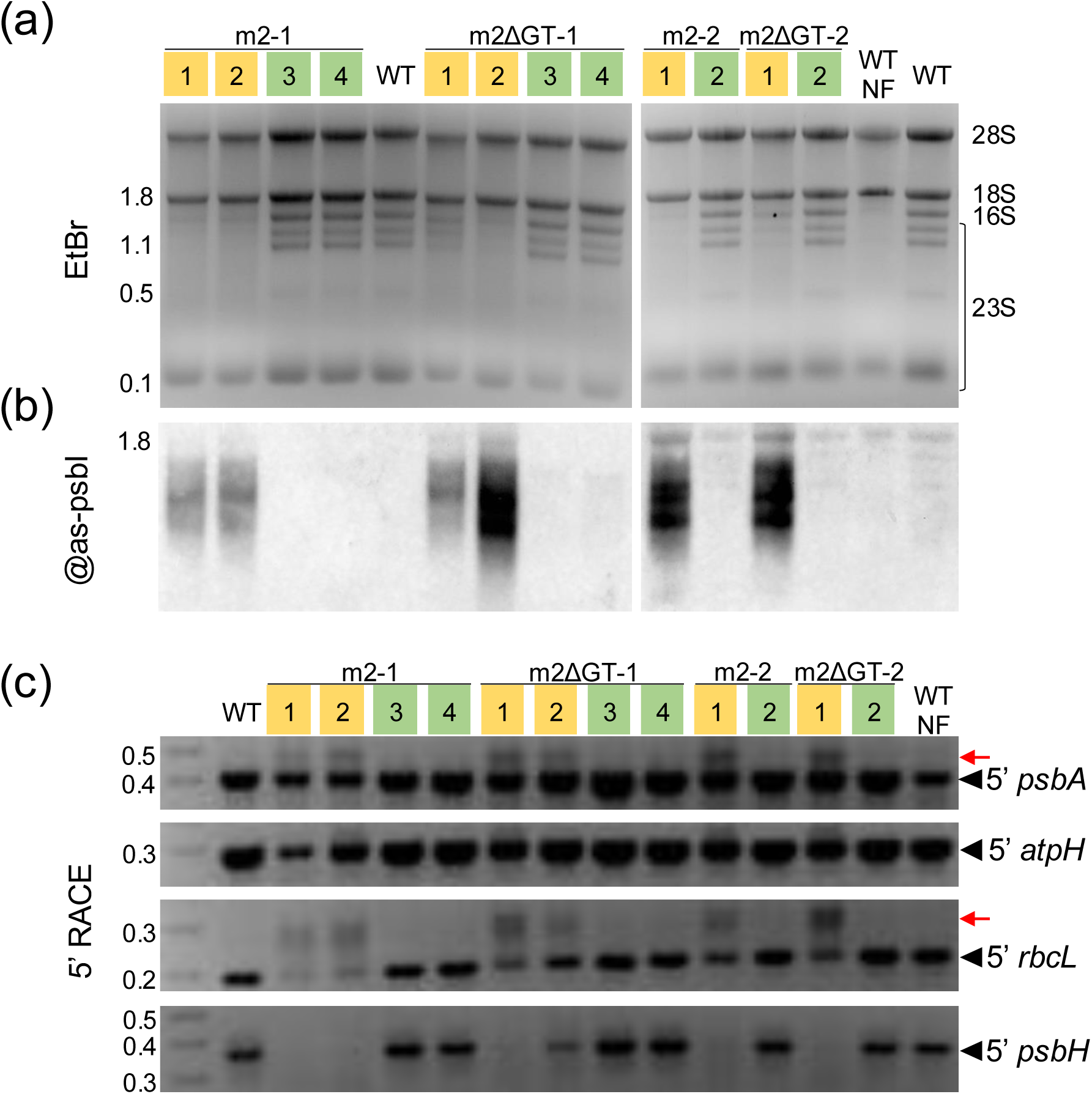
Chlorotic plants have RNA quality control defects. Green and yellow boxes indicate green and chlorotic plants, respectively. WT NF indicates sample from WT plants grown on media containing 5 µM Norflurazon for 7 days. Sizes in kb are shown to the left of gels. (a) Ethidium bromide-stained gel of 4 µg total RNA from the indicated genotypes. Major rRNA species are indicated on the right, with 16S/23S being chloroplastic and 28S/18S mainly cytosolic. (b) RNA gel blot detection of *psbI* antisense transcription by hybridization with a biotinylated oligonucleotide, from the same samples as in panel (A). (c) 5’ RACE analysis of *psbA*, *atpH*, *rbcL* and *psbH*. 5’ termini of indicated transcripts were amplified with gene-specific and adapter-specific primers from the same samples as in panel (A). Black arrowheads indicate the major 5’ termini. Red arrows indicate the 5’ extended species.

RNJ is also involved in 5’ end maturation, as originally suggested by studies of 5’ end resection eventually impeded by PPR proteins (Pfalz *et al*., 2009). When VIGS was used to reduce *RNJ* expression in tobacco, 5’-extended RNA species were identified for *psbA*, *atpH*, *rbcL* and *petB* (Luro *et al*., 2013), corresponding to known targets of 5’ exonucleolytic processing. We employed 5’ RACE to evaluate 5’ maturation of the first three of these transcripts, plus *psbH*, comparing green and chlorotic plants (Figure 5c). In the chlorotic plants, in addition to the major 5’ terminus, discrete 5’-extended transcripts were detected for *psbA* and *rbcL* (red arrows in Figure 5c). The major 5’ end of *psbH*, however, was barely detectable, potentially indicating a broader defect in the complex maturation of the *psbB* polycistron (Stoppel and Meurer, 2012). In contrast, results for the 5’ end of *atpH* were inconsistent, with four of six RNA samples from chlorotic plants appearing to harbor a minor 5’-extended species, but the other two resembling the green plant results (Figure 5c).

To confirm that these results were not due to a pleiotropic effect of chlorosis, we treated WT plants with Norflurazon, an inhibitor of phytoene desaturase, to chemically induce chlorosis. Although like the chlorotic plants studied here Norflurazon-treated plants lack plastidial rRNA (Figure 5a, lane WT-NF), they do not accumulate *psbI* asRNA or any extended 5’ termini. This result strongly suggests that RNA quality control defects observed in the *motif II* chlorotic plants is somehow related to expression of the RNJ construct.

### Chlorotic plants that fail to accumulate transgene protein continue to express both endogenous and transgene RNJ genes

The fact that only homozygous transgenic lines developed chlorosis suggested a dominant effect driven by gene copy number. Because the functional RNJ form is a dimer or tetramer (Newman *et al*., 2011; Zhao *et al*., 2015; Zheng *et al*., 2017; Raj *et al*., 2020; Bao *et al*., 2023; Tejada-Arranz *et al*., 2023), we speculated that the emergence of chlorosis might be triggered by incorporation of the catalytically inactive *motif II* protein into heterooligomers containing WT RNJ, forming a low/null activity complex leading to effects reminiscent of *RNJ* silenced plants. Because this mechanism would rely on the presence of the mutant protein, we used immunoblots to detect the transgenic protein through its Flag tag. In contrast to what this hypothesis would predict, m2 and m2ΔGT proteins were readily detected in green plants of the four lines but were undetectable in all chlorotic plants (Figure 6). Therefore, poisoning of RNJ oligomers cannot explain the results presented to this point.

**Figure 6.**
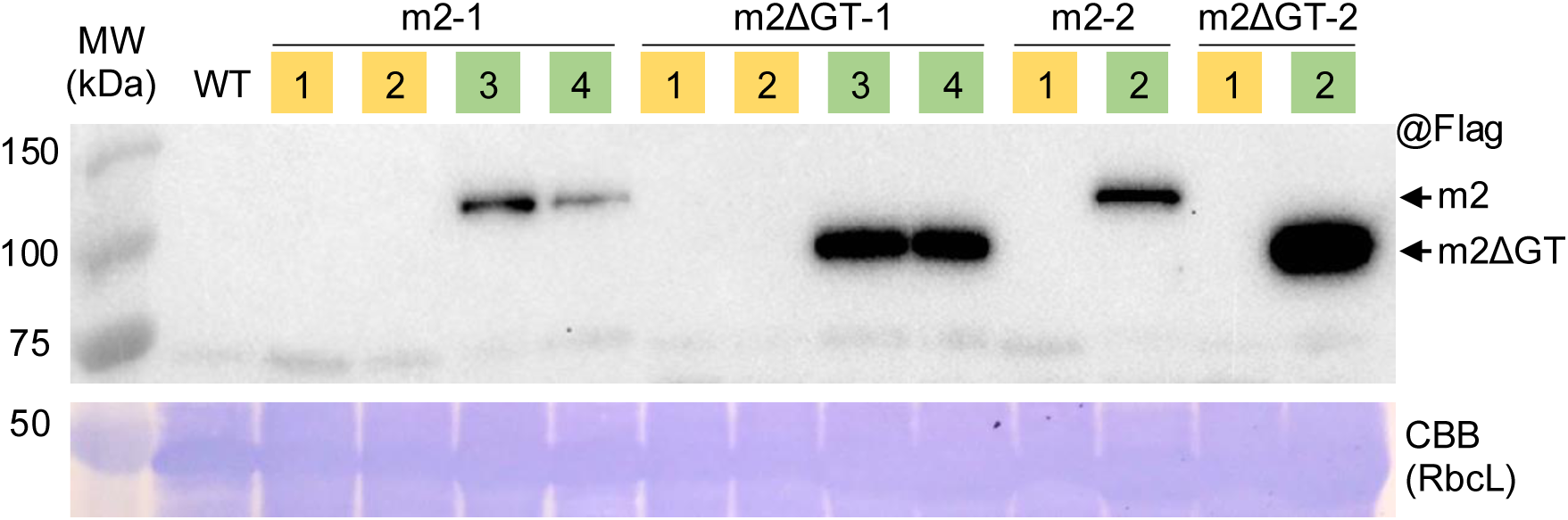
Chlorotic plants do not accumulate transgene-encoded protein. Protein analysis of indicated genotypes, with green and yellow boxes indicating green and chlorotic plants, respectively. Top, immunoblot with anti-Flag antibody. Lower, Coomassie-stained gel cropped to show the region of the Rubisco large subunit.

The next logical explanation for an *rnj*-like phenotype would be an absence rather than a poisoning of the RNJ complex. While we could not evaluate directly the abundance of the native RNJ protein due to the lack of a suitable antibody, transcript accumulation was evaluated by RT-qPCR, using the same primers as for qPCR to distinguish endogenous and transgene expression. Both the endogenous and transgenic *RNJ* mRNAs accumulate in the chlorotic m2-1 and m2ΔGT-1 plants, as well as in green plants (green and blue bars, respectively, in Figure 7). The measured level of endogenous *RNJ* mRNA varies from 50-100% of the WT control across lines and plant phenotypes and does not differ between samples. There is some suggestion that transgene mRNA has lower accumulation in chlorotic vs. green plants of a given line, although this could not be statistically supported. In conclusion, transcriptional silencing of either endogenous or transgenic *RNJ* does not explain the observed phenotype. Given the disappearance of transgenic RNJ in chlorotic plants, we propose that post-transcriptional gene silencing (PTGS) results in the lack of synthesis/degradation of both endogenous and transgenic RNJ proteins.

**Figure 7.**
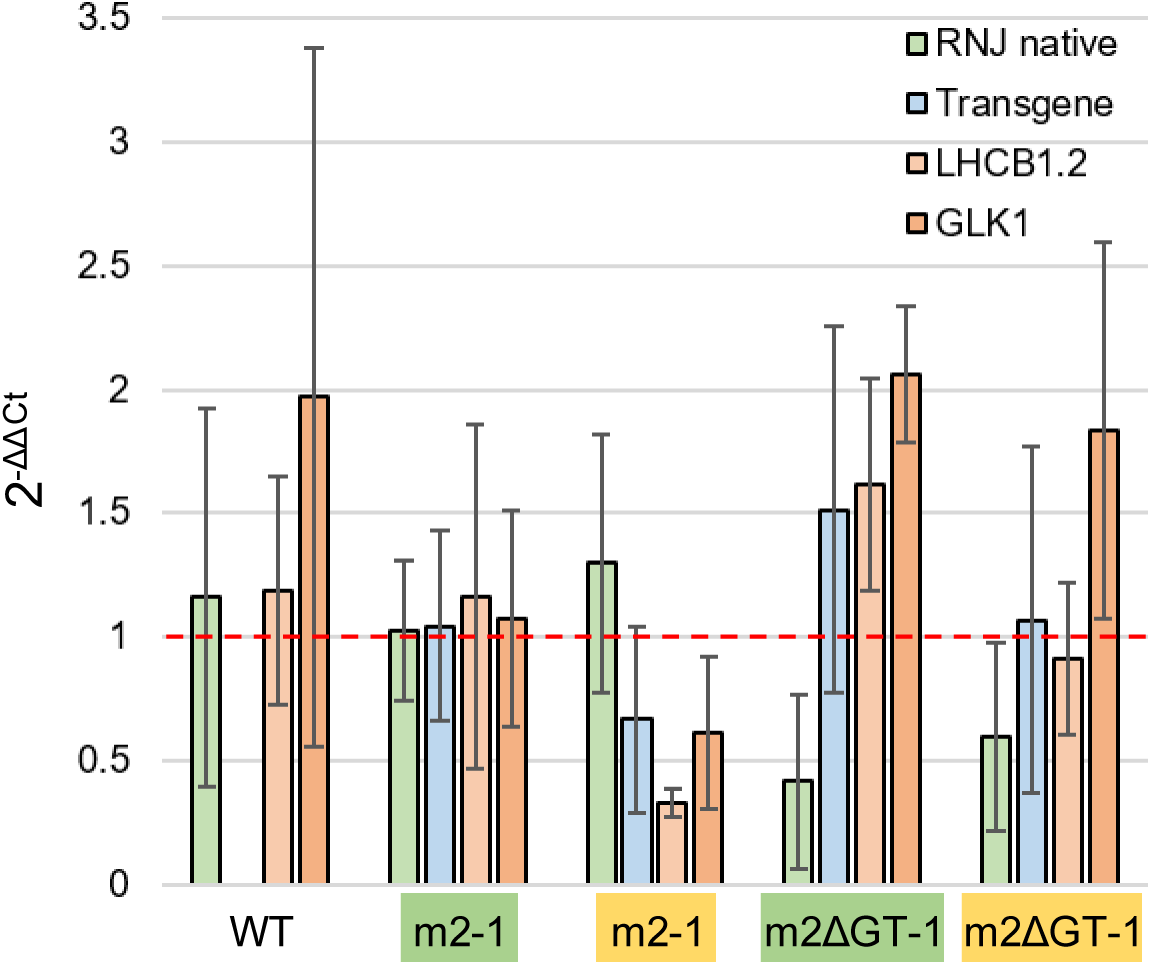
*RNJ*, transgene, *LHCB1.2,* and *GLK1* expression in green and chlorotic plants. Green and yellow boxes indicate green and chlorotic plants, respectively. Quantitative RT-PCR was used to query expression levels of four genes in the five genotypes indicated. RNJ native, the endogenous *RNJ* gene. Transgene, the respective transgene present in the *motif II* lines. Expression was normalized against m2-1 green plants and relative expression is calculated using *ACTIN1* and *UBC21* as reference genes. Values are mean ± SD (n=4).

Because PTGS can arise from retrograde signaling pathways, we checked mRNA levels for two genes, *LHCB1.2* and *GLK1*, two photosynthesis-associated nuclear genes (PhANGs) that were shown by others to be suppressed by silencing of *RNJ* in tobacco (Ganusova *et al*., 2020). In line m2-1, accumulation of both markers was reduced by about 50%, similar to the results reported by Ganusova *et al*. This effect was similar for *LHCB1.2* in m2ΔGT-1, however average *GLK1* mRNA accumulation did not change, although expression was highly variable in m2ΔGT-1 chlorotic plants as can be seen from the standard deviation. Overall, RT-qPCR results are partly consistent with induction of a previously studied RNJ-related retrograde signaling pathway, one that in this case results in diminution of RNJ activity through a post-transcriptional mechanism.

### Plants undergoing transgenic RNJ silencing do so heterogeneously

At the end point, chlorotic and green plants differ in their plastid and nuclear transcriptomes, and in transgenic protein accumulation. At an early stage of development, however, all seedlings are green. This raises the question of what triggers chlorosis in a subpopulation of genetically identical plants. To begin addressing this question, we qualitatively investigated selected molecular phenotypes of plants fated to develop chlorosis. To do so we used line m2ΔGT-1, which generates chlorotic plants that are frequent and easy to identify, although we suspect results would be similar for the other transgenic lines.

The experiment began after germination and transfer to soil, when seedlings were genotyped for transgene copy number, thereby identifying those with the potential to develop chlorosis. Samples were collected from transgene homozygotes before chlorosis could be detected (pre), then one (+1) and ten (+10) days after the initial appearance of chlorosis. As controls, samples were collected from hemizygous seedlings of a similar age and developmental stage (Figure 8a).

**Figure 8.**
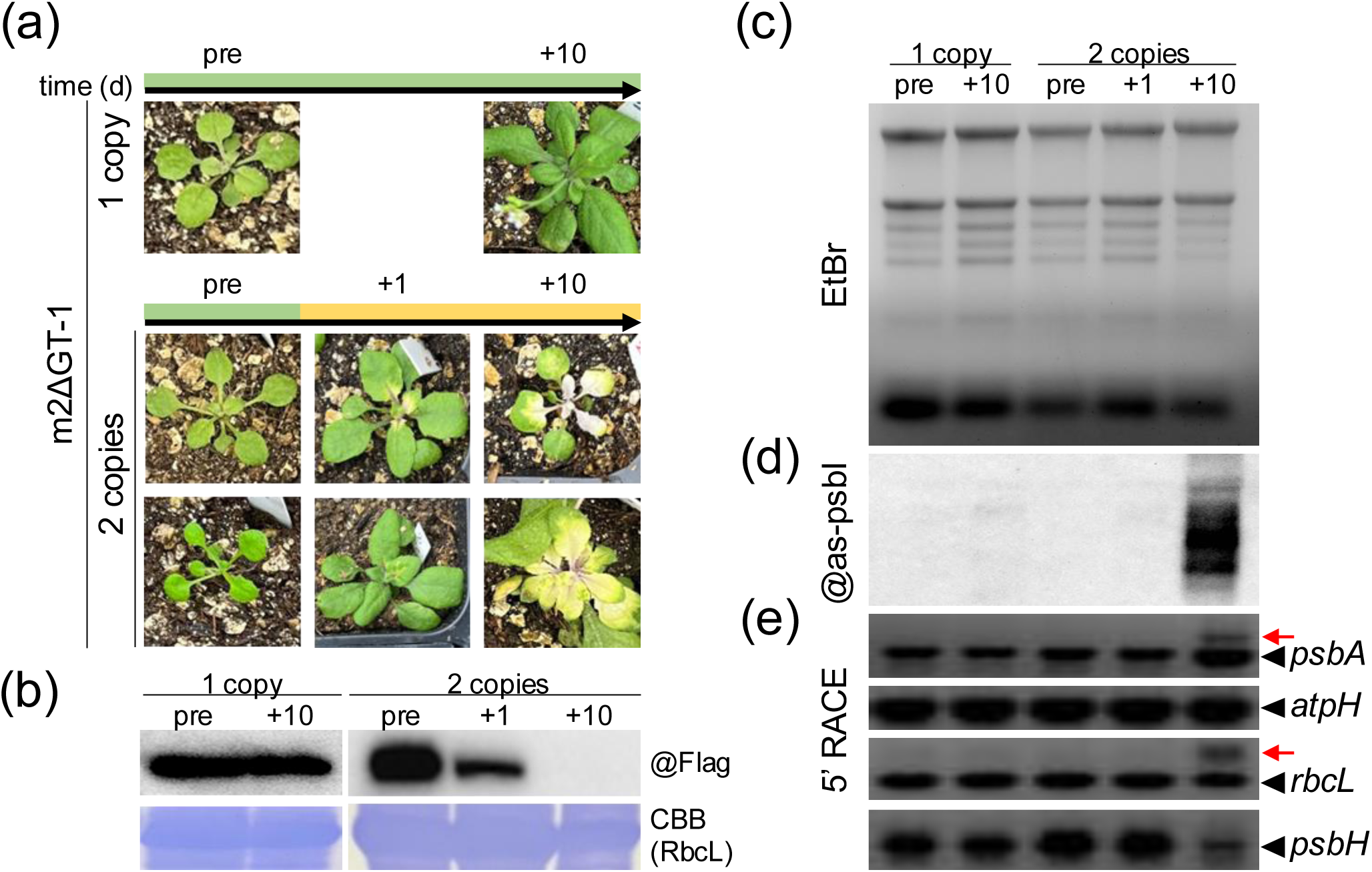
Time course of the molecular phenotype of m2ΔGT-1 plants during chlorosis development. (a) Images of representative plants containing one copy of the transgene at the pre and 10 d time points (top). Images were also obtained for plants containing two copies of the transgene before chlorosis appears (pre), then one and 10 days after its appearance (bottom). (b) Protein analysis of samples according to the design depicted in panel (a). Top, immunoblot with anti-Flag antibody showing presence/absence of the m2ΔGT protein. Lower, Coomassie-stained gel cropped to show the region of the Rubisco large subunit. (c) Ethidium bromide-stained gel. (d) RNA gel blot from the same samples as in (c). (e) 5’ RACE analysis from the same samples as in panel (c).

We first checked the accumulation of transgenic RNJ protein using immunoblots, as shown in Figure 8b. Pre-chlorosis, transgenic RNJ was readily detected in all samples. The signal decreased markedly as soon as chlorosis developed (+1), and was undetectable at +10 for homozygous, chlorotic plants, as shown earlier (Figure 6). No effect on RNJ accumulation was observed in hemizygous plants, which remained green. To investigate the time course of protein disappearance more finely, individual homozygous plants were sampled pre-chlorosis, the day the chlorosis appeared (+0) and three days later (+3). Although for all plants transgenic RNJ accumulated pre-chlorosis and decreased thereafter, there was variation at each time point (Figure S4). The results for “+0” samples suggest that a few of the “pre-chlorosis” plants were in fact already induced for the response leading to chlorosis. In subsequent days, we can imagine that as a cell-autonomous phenomenon is in transition to a systemic one, RNJ accumulation is varying as the sum of a heterogenous collection of cell types. In addition, whatever the signal is, it likely varies in strength and thus effect from plant to plant at this early stage.

To probe this concept of heterogeneity in more depth, representative homozygous plants from pre, +0, +3, +5 and +10 time points were dissected, and each leaf or similar group of leaf sections was examined individually for transgenic RNJ accumulation. Figure S5 shows that all samples from a “pre” plant accumulated transgenic RNJ, as expected. On day 0, there was substantial leaf-to-leaf variability, where transgenic RNJ accumulation could largely be associated with the intensity of chlorosis. At day +3, fewer samples had detectable signals, and these were uniquely in green tissue. By day +5, even green leaves only had a faint signal. At the end point (>+10), green leaves did not accumulate transgenic RNJ, suggesting that a systemic silencing had occurred. An interesting exception at day +10 was the “m” samples, referring to the rosette “leftovers” after excising all leaves. Here, a signal persisted, suggesting that silencing was restricted to developed leaves. Curiously, this “m” signal was not observed earlier at +0, +3 or +5.

In contrast to the heterogenous and progressive loss of RNJ, transcriptome changes only appeared later in chlorosis. For example, a decrease in rRNA became visible only at +10 (Figure 8c), as was the case for detection of *psbI* asRNA (Figure 8d) and 5’ end extensions (Figure 8e). Taken together, our results show that before the appearance of chlorosis, plants with two copies of the transgene are indistinguishable from those with one copy for the parameters measured. Decreased accumulation of transgenic RNJ is an early event associated with chlorosis (between +0 and +3) and later becomes systemic, whereas RNA phenotypes are trailing and presumably signal the suppression of WT RNJ at the protein level, which may or may not fully parallel the decrease of the inactive, tagged version. Whether RNA phenotypes occur earlier in individual cells or sectors, or only at +10, cannot be determined from our experiments.

### Attempts to overexpress active RNJ result in similar and even stronger phenotypes

The data presented above do not support a dominant negative model for transgenic RNJ carrying motif II mutations in terms of inactivating endogenous RNJ, but a defective RNJ protein might nonetheless be required to trigger the observed responses. To test this hypothesis, we created a construct designed to overexpress active RNJ (hereafter called WTox). This construct is identical to m2 apart from encoding a WT rather than mutant version of motif II. In other words, it is the same construct earlier used to complement the *rnj* mutant (WTc; Baudry *et al*., 2026), except that the 35S promoter instead of the native *RNJ* promoter drives its expression (Figure 9a).

**Figure 9.**
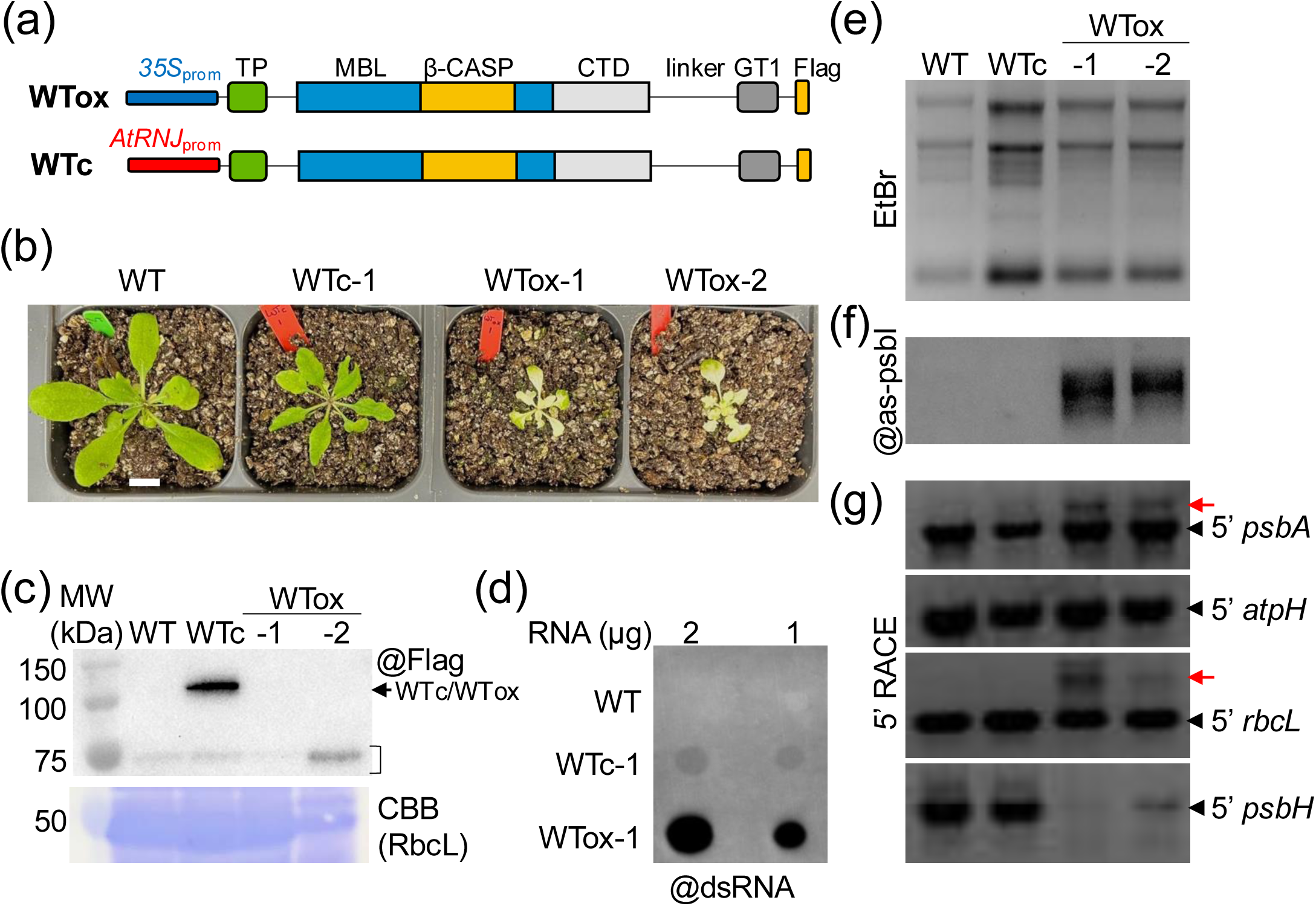
Characterization of plants overexpressing active RNJ. (a) Constructs used to transform the heterozygous *rnj-2* mutant. The CaMV 35S promoter (WTox) or native *RNJ* promotor (WTc) was used to drive expression of wild-type AtRNJ, flanked by a Flag epitope tag. (b) Untransformed WT, a representative WTc-1 plant, and two independent events for the WTox construct. Plants were grown in long day conditions at 22°C (scale bar = 1 cm). (c) Protein analysis of the indicated genotypes. (d) dsRNA detection by dot blot analysis for the indicated lines. (e) Ethidium bromide-stained gel of 4 µg total RNA from the indicated genotypes. (f) RNA gel blot detection of *as-psbI* from the same samples as in panel (e). (g) 5’ RACE analysis of the same samples as in panel (e).

As was done for constructs expressing motif II mutations, we transformed heterozygous *rnj-2/+* plants and screened for transformation events. These turned out to be very rare, and after extensive screening for hygromycin resistant seedlings, only two T1 events were obtained from ten transformed T0 plants, an efficiency at least 20-fold lower than with the motif II constructs. Those T1 plants quickly developed severe chlorosis; however, a few seeds could be obtained. T2 seedlings were green at the cotyledon stage, but 100% of the examined seedlings were spindly and developed severe chlorosis (Figure 9b), which began to appear sooner than in *motif II* lines, as early as one week DAS (Figure S6a). None of the T2 plants was identified as homozygous for the *rnj-2* T-DNA insertion, and all of them were hemizygous for the transgene (Figure S6b). These results suggest that this construct is unable to complement background embryo lethality, and that *RNJ/RNJ* or *rnj/RNJ* seeds possessing two transgene copies are unable to germinate.

We examined the RNA phenotypes of WTox lines in comparison to WTc-1, which is a phenotypically normal if somewhat smaller plant than a true WT, and expresses the same active full-length enzyme as WTox but under control of the *RNJ* native promotor (Baudry *et al*., 2026). In contrast to WTc and *motif II* transgenics at a green stage and similarly to the chlorotic plants described above, chlorotic WTox plants do not accumulate the transgenic protein (Figure 9c). They do, however, accumulate dsRNA (Figure 9d), as well as the diagnostic RNA markers characteristic of RNJ deficiency presented earlier for *motif II* plants (Figures 9e-g). When sampled before chlorosis, WTox plants do accumulate transgenic protein and the chloroplast transcriptome appears to be normal (Figure S7). Therefore, the effects of expressing WTox and m2/m2ΔGT are similar but are accelerated, and occur at a lower transgene ploidy for WTox.

Altogether, our results show that overexpressing either active or inactive versions of RNJ can lead to chlorosis and chloroplast RNA quality control deficiency, supporting our earlier conclusion that the observed phenotypes of *motif II* lines are unlikely to be explained by inactivity of the enzyme. In addition, because 100% of WTox seeds viable on hygromycin medium have only one copy of the transgene and develop severe chlorosis, it suggests that expressing active RNJ under the CaMV 35S promotor is more deleterious than expressing the inactive version. Possible mechanisms underlying this phenomenon or related but different phenomena for *motif II* vs WTox results, are discussed below.

## DISCUSSION

### Overexpression of RNase J is deleterious to plant growth

In an approach designed to trap RNJ substrates through expression of a catalytically inactive enzyme, we expressed two versions of RNJ under the CaMV 35S promoter. This resulted in chlorosis and severe defects in plastid gene expression, a phenotype that could be replicated by 35S-driven expression of wild-type RNJ. Our observations are in stark contrast with the few prior examples of overexpression of chloroplast ribonucleases in Arabidopsis. In one study, a mutant for polynucleotide phosphorylase was complemented using a 35S-driven construct (Walter *et al*., 2002). An up to 20-fold increase in PNPase content was observed in some lines without a deleterious effect, whereas several lines were also obtained that were postulated to abolish PNPase expression through cosuppression. It has been shown that cpPNPase null mutants have mild effects on Arabidopsis development, such as transient chlorosis (Germain *et al*., 2011), in spite of major effects on the chloroplast transcriptome (Castandet *et al*., 2013). A mutant of RNR1, which is a protein in the RNase II family, was also complemented with a 35S promoter-driven construct (Perrin *et al*., 2004). While Perrin *et al*. focused on RNR1 as a mitochondrial protein, later work showed that it is also targeted to chloroplasts and involved in 3’ end maturation and RNA decay (Germain *et al*., 2012; Bollenbach *et al*., 2005). No adverse effects of RNR1 complementation by the 35S constructs were noted. Finally, 35S-driven complementation of a mutant lacking CSP41b, one of the two paralogous chloroplast endonucleases, generated plants at least as robust as the WT (Qi *et al*., 2011). The other major chloroplast RNase is RNase E, whose absence strongly affects mRNA patterns from gene clusters and causes seedling chlorosis (Walter *et al*., 2010). To our knowledge, overexpression of RNE has not been attempted. Taken together, this literature suggests that RNJ is particularly sensitive to increased expression as compared to other chloroplast ribonucleases.

Two previous studies demonstrated complementation of the *rnj* mutant. In one study, a genomic fragment of the *RNJ* promoter and coding region was used to complement three different T-DNA insertion mutants successfully (Chen *et al*., 2015). Only the embryo phenotypes of these transformants were reported, so whether the plants were phenotypically indistinguishable from the WT, or whether any chlorotic progeny were observed, is unknown. Our own study used a cDNA rather than a genomic *RNJ* transgene to complement one of the T-DNA mutants; however, the same promoter was used as in Chen *et al*. (2015). We observed that plants complemented with WT RNJ were smaller but otherwise resembled true WT plants (Baudry *et al*., 2026). If excess RNJ in any form is deleterious to plant growth, it is possible that our WTc line being smaller is due to a higher than ideal abundance of RNJ, consistent with the observed higher expression of WTc mRNA compared to endogenous *RNJ* reported in our previous study, although there are other possible explanations (see discussion in Baudry *et al*., 2026).

Here, the overexpression was attempted using 35S-driven transgenes that tend to be highly expressed in *Arabidopsis* cotyledons and young leaves (Kiselev *et al*., 2021). These are the same tissues in which chlorosis arose in our experiments and in which RNJ-GUS fusions suggest RNJ is also highly expressed and therefore likely to be of most importance (Chen *et al*., 2015). Additionally, at least the motif II mutated transgenic proteins were easily detected in non-chlorotic tissues (Figure 6), consistent with robust expression. We therefore speculate that promoter strength (meaning the 35S promoter and 5’ UTR) is a driving factor in the phenomenon under study.

### Overexpression of transgenic RNase J silences both transgenic and endogenous enzymes

The RNA phenotypes we observed in chlorotic plants appeared to be a phenocopy of those we previously reported for VIGS-silenced plants (Luro *et al*., 2013; Sharwood *et al*., 2011), with the exception that we did not observe any effect on the *atpH* 5’ end (Figures 5c and 8e). Apart from this minor distinction, the accumulation of *psbI* asRNA, diminution of chloroplast rRNA, and appearance of other 5’ end extensions, were all found in the VIGS plants, along with generalized dsRNA accumulation. These extensive similarities to VIGS plants strongly suggest that although we cannot detect the WT RNJ protein directly, it is undergoing the same fate of silencing as the transgenic proteins. In other words, our attempts to overexpress RNJ have resulted in silencing of both the transgenic and endogenous enzymes.

Cosuppression pathways could be an obvious explanation of the phenomenon observed here. It is common to trigger post-transcriptional gene silencing via overexpression of transgenes and indeed, in addition to the experiment with PNPase mentioned above, 35S promoter-driven constructs were purposefully used to silence embryo-defective genes in *Arabidopsis*, including several encoding pentatricopeptide repeat proteins involved in chloroplast gene expression (Chen *et al*., 2020). Further analysis showed that the transgenes acted in an RDR6-dependent manner to suppress mRNA accumulation for the targeted genes. Moreover, it has also been shown in Arabidopsis that the AGO1-dependent pathway can involve translational repression along with RNA degradation (Lanet *et al*., 2009). Our results, however, are unlikely to be explained fully by these mechanisms as neither transgene nor endogenous *RNJ* mRNA accumulation changes could account for observed phenotypes (Figure 7). Although we observe partial reduction of transgene mRNA, our RT-qPCR used samples that detailed analysis showed were likely to be phenotypic mosaics (Figure S5), making it impossible to know whether there was substantial suppression of *RNJ* transgene mRNA accumulation. Nonetheless, expression of the endogenous *RNJ* mRNA stayed the same, which is inconsistent with a cosuppression hypothesis involving RNA degradation. This strongly suggests that RNJ abundance is decreasing either through proteolysis alone or through a combination of proteolysis and translational repression.

### Mechanisms leading to RNJ inhibition

If one accepts the premise that attempted overexpression of RNase J causes the observed phenotypes outside of a previously described cosuppression pathway, what could be the mechanism(s) explaining the eventual deficiency of the RNJ protein? Answers to this question must account for the fact that both active and inactive RNJ proteins can trigger the response, and that 35S-driven expression of the WT protein (WTox lines) is a far more efficient trigger. A model that frames the following discussion is depicted in Figure 10. It assumes that downstream events are triggered by “excessive” RNJ, accounting for the absolute dependence on motif II mutant transgene homozygosity for the development of chlorosis. We further assume that the abundance of endogenous RNJ is not affected by zygosity at the *RNJ* locus. This is based on the facts that 1) *RNJ/RNJ* and *rnj/RNJ* plants are equally prone to chlorosis; and 2) there is no evidence that *rnj/RNJ* plants lacking transgenes are distinguishable at the molecular level, i.e. there is no evidence for haploinsufficiency.

**Figure 10.**
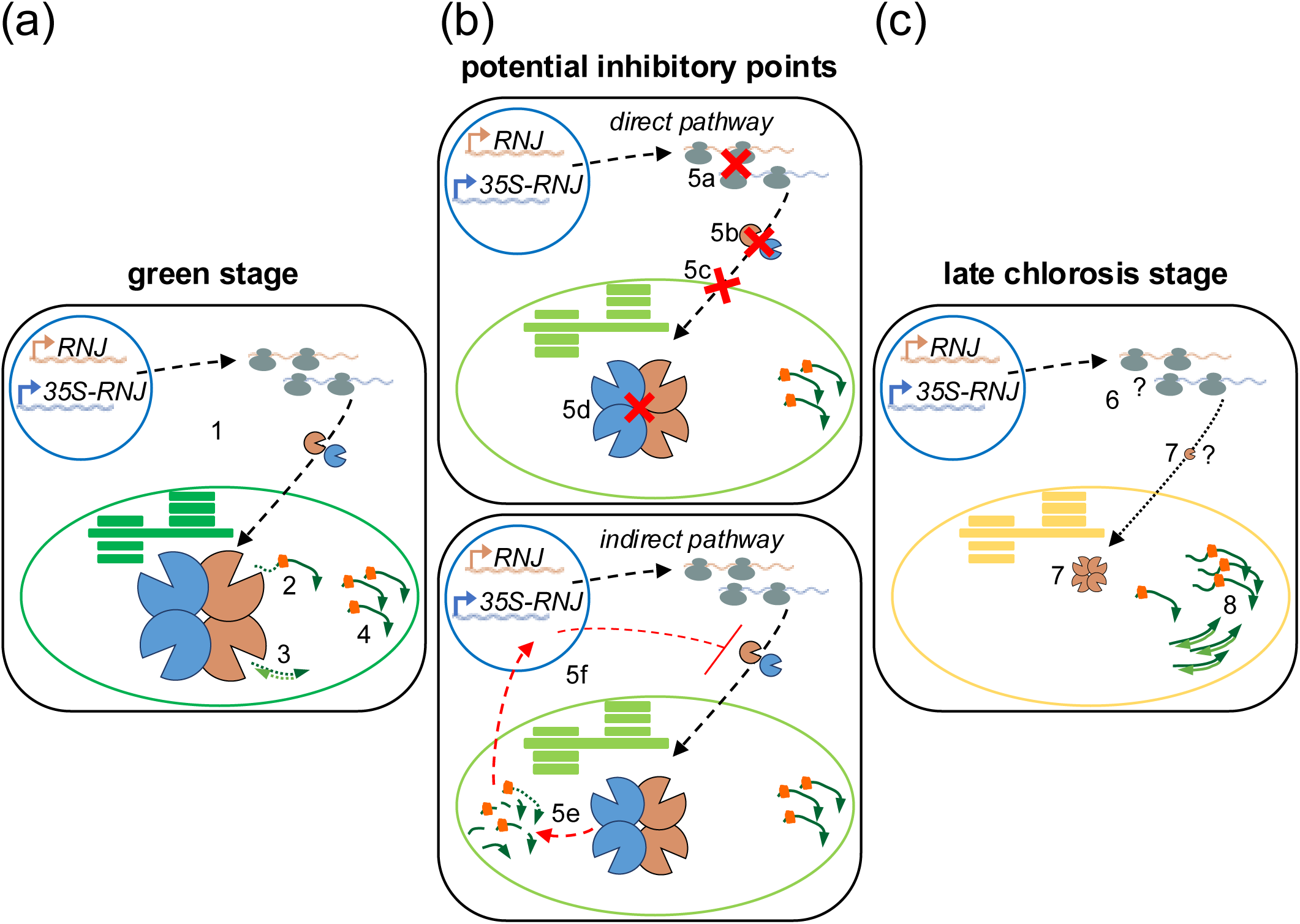
Model depicting the path to RNJ inhibition in plants developing chlorosis. The three columns from left to right illustrate RNJ-related mechanisms leading to RNJ inhibition in homozygous transgenic plants that will ultimately develop chlorosis. Not shown are WT plants or transgenic plants with genotypes that do not prime the plants for chlorosis, namely hemizygous m2 or m2ΔGT plants. The diagrams also do not differentiate between active ( WTox) and inactive (m2/m2ΔGT) forms of RNJ: both are depicted in blue. For simplicity, the many possible homo- and heterooligomers are not shown: a simple 2:2 tetramer is used to illustrate situations where both transgenic and endogenous forms are present. (a) In green plants pre-chlorosis, both endogenous RNJ (brown) and transgenic RNJ (blue) are expressed and imported into the chloroplast (1) forming the RNJ complex, which is hypothesized to be present in excess, as indicated by the large size of the tetramer. Active subunits of these complexes process 5’ transcript termini (2) and degrade asRNA when present in dsRNA duplexes (3), allowing the accumulation of only 5’-matured sense RNAs (4) whose termini are protected by RNA-binding proteins (orange squares). (b) As a threshold of excessive RNJ activity is detected in a chloroplast or cell, a mechanism leading to RNJ inhibition is initiated and early chlorosis is observed. The top panel represents a direct mechanism that could involve inhibition of RNJ translation (5a), cytoplasmic proteolysis (5b), restrictions on import (5c), and/or RNJ proteolysis within the chloroplast (5d). The bottom panel shows an alternative indirect pathway in which excessive RNJ activity overprocesses sense transcripts (5e), triggering the RNJ inhibition mechanism diagrammed in the top panel through retrograde signaling (5f). At the stage depicted in column (b) the quantity of RNJ has begun to decrease, however RNJ transcripts and protein are still present and thus sufficient RNJ activity for transcript processing. (c) At later stages of chlorosis, RNJ transcripts are still present, with the question mark indicating that their translational status is unknown (6). The transgene - encoded protein is absent, and the native RNJ protein is hypothesized to be of very low abundance, or absent (7). As a result of diminished RNJ activity, 5’-extended (unprocessed) transcripts and asRNA (in form of dsRNA) accumulate (8).

One mechanistic possibility is that the overexpression of RNJ leads to its accumulation in an unprocessed and/or misfolded form, either in the cytoplasm (Figure 10b, point 5b), in transit across the chloroplast envelope (point 5c), and/or within the chloroplast (point 5d). The forced accumulation of unprocessed chloroplast precursor proteins in the cytoplasm is known to inhibit chloroplast development and to trigger the ubiquitin-proteasome system to degrade the mis-processed protein (Hong *et al*., 2025). While this could potentially explain the disappearance of RNase J, we never detected the accumulation of the precursor protein and the ability of the WTc construct to complement the homozygous *rnj* mutant shows that the transgenic protein is correctly imported into to the chloroplast (Baudry *et al*., 2026). A more likely possibility is that the enzyme is degraded within the chloroplast stroma by its proteostasis machinery. In particular, the Clp protease complex degrades aggregated or misprocessed proteins and has been linked to a previous failed attempt to overproduce the chloroplast ribosomal protein PRPS1, which like RNJ is required for embryo development in Arabidopsis (Tadini *et al*., 2023). In general, there are multiple triggers and pathways that the chloroplast uses for proteostasis, one or more of which could be at play in our situation (reviewed in Jeran *et al*., 2025).

A subsidiary consideration is that our transgenic plants are green before chlorosis may initiate, meaning that overall RNJ activity is at first within the range required to ensure correct chloroplast development. This suggests that overabundance occurs later, or that an overabundant level of RNJ is tolerated at early stages, but at some development stage begins to exert a negative effect on chloroplast gene expression (point 5e) that secondarily leads to its inhibition through signaling (point 5f). In molecular terms, given that RNJ is not a sequence-specific enzyme, we posit that above a certain threshold it begins to degrade sense mRNAs and/or rRNAs that are normally protected by secondary structures and/or RNA-binding proteins. Indirect evidence for this is the appearance of new degradation products in longer incubations of AtRNJ with *in vitro* substrates (Halpert *et al*., 2019). Similarly, when *B. subtilis* RNJ was used as a probe for RNA secondary structures, over time additional cleavages occurred in normally refractory regions (Daou-Chabo and Condon, 2009). The 3’→5’ exonucleases RNase II and RNase R also have concentration-dependent activities, among others (Cheng and Deutscher, 2002). Thus, increased RNJ activity in WTox could lead to more undesirable RNA degradation as secondary structures breathe or as RNA-binding protein sites are either initially formed through transcription or periodically vacated.

How increased abundance of mutant RNJ subunits might eventually trigger the same effect, if less efficiently, is somewhat puzzling. A possibility is that motif II mutant proteins do not impede the activity of co-assembled WT subunits but instead increase the overall activity of RNJ by allowing the assembly of additional stable tetramers. In other words, the model proposes that RNJ abundance is usually capped (hence the similarity of *RNJ/RNJ* and *rnj/RNJ* plants) and that the presence of excess protein enabled by a transgene allows this cap to be breached. While we cannot directly detect the WT protein, we assume that motif II proteins can assemble with the WT version, since only the catalytic site is mutated. To our knowledge, this has not been tested *in vitro*, although it was shown previously that paralogous BsRNJ1 and BsRNJ2 form heterotetramers (Mathy *et al*., 2010).

It is likely that retrograde signaling takes place as either a direct or indirect outcome of altered RNJ activity. The nature of chloroplast molecular signals that trigger retrograde signaling is a highly active area of study and there is considerable literature linking the state of plastid gene expression to retrograde signaling in a pathway involving *GUN1* (Jan *et al*., 2022). In our experiments, roughly 25% of genetically eligible plants underwent chlorosis for motif II mutants, whereas 100% of WTox plants became chlorotic (Figure 2b and S6a). Thus, the threshold for dysfunction is invariably met in WTox plants, whereas it is only sometimes reached in *motif II* lines. This is consistent with any of the models proposing that high expression of mutant RNJ leads indirectly to transcriptome defects by allowing greater accumulation of endogenously encoded active RNJ, whereas WTox increases overall RNJ activity directly.

## MATERIALS AND METHODS

### Plant materials and growth conditions

The T-DNA insertion mutant *rnj-2* (SAIL_343_B07), used as the genetic background, was previously described (Chen *et al*., 2015). All plants, on media or on soil, were grown in long day conditions (16h light/ 8h dark) at 22°C under fluorescent lighting (120–140 μmol.m^−2^.s ^−1^). Seeds were sown on 1/2 MS medium containing 33 µg/mL hygromycin B (Phyto Tech Labs), 2–3 week old seedlings were transferred to soil. To monitor the development of chlorosis, groups of 88 seeds were sown on media and kept in the dark at 4°C for 48 h before being placed in the light. Plant development and coloration was noted daily. The proportion of chlorotic plants was expressed as a percentage of hygromycin resistant (transgene-containing) plants.

### Cloning, plant transformation, and selection

Sequences encoding RNJ *motif II* mutants flanked by Gateway *att*B recombination sites were synthesized by Twist BioScience (San Francisco, USA). m2 corresponds to the full-length Arabidopsis RNJ CDS with no intron and carries the same mutated motif II (H177A, D179K, H180A) as in Halpert *et al*. (2019). m2ΔGT corresponds to the m2 construct but the GT1 domain is truncated (amino acids 805 to 904 are removed) as previously described (Halpert *et al*., 2019; Baudry *et al*., 2026). WTox corresponds to m2 in which the motif II mutations were reverted to WT by point mutation amplification. ORFs were inserted into pDONR221 using Gateway BP clonase (Invitrogen), then subcloned into pGWB511 using Gateway LR clonase (Invitrogen), which allows expression from the CaMV 35S promoter and adds a C-terminal FLAG tag. Heterozygous *rnj*-2/+ plants were transformed using C58C1 pMP90 *Agrobacterium tumefaciens* by floral dip (Clough and Bent, 1998) and selected on 1/2 MS media containing 33 μg/mL hygromycin B. Line WTc-1 was previously characterized (Baudry *et al*., 2026) and corresponds to WT full-length RNJ driven by its native 2.0 kb promoter. Primers used for genotyping are given in Table S1.

### Quantitative PCR

Transgene copy number was estimated by qPCR on gDNA using iTaq-Univeral SYBR Green Supermix (Bio-Rad). For each plant, the mean of two technical replicates of transgene signal was normalized by the mean of the native *RNJ*. Primers used are listed in Table S1.

### RNA extraction, RNA gel blots, 5’ RACE, cDNA synthesis, RT-qPCR, and dot blots

Leaf total RNA was extracted using the NucleoZOL protocol (Macherey-Nagel) and RNA quality was checked by analyzing 4 μg in a 1.2% formaldehyde-agarose gel. For gel blots, RNAs were blotted overnight from formaldehyde-agarose gel onto Hybond-N+ (GE Healthcare) in 25 mM sodium phosphate pH 6.8 buffer. Blots were probed as described (Baudry *et al*., 2022) with the following minor modifications. Hybridization was performed with biotinylated oligonucleotides at 48 nM, which had been denatured at 70°C for 10 min. Blots were washed twice 10 min in 3X wash buffer (3X SSC, 20 mM Na_2_HPO_4_ pH7.5, 5% SDS). For 5′ RACE, an RNA adapter was ligated to 2 μg RNA using T4 RNA ligase (NEB) as described (Luro *et al*., 2013). cDNA was synthesized using SuperScript III (Invitrogen). 5′ termini were amplified with GoTaq (Promega), using an adapter-specific primer and a gene-specific primer. RT-PCR products were separated in 2% agarose gels. Gene expression of transgene, native *RNJ* (At5g63420), *LHCB1.2* (At1g29910), *GLK1* (At2g20570), was measured by RT-qPCR, performed as for qPCR. Four biological replicates were obtained and for each the mean of two technical RT-qPCR replicates were normalized with the mean of the reference gene *ACTIN1* (At2g37620) and *UBC21* (At5g25760) using the ΔΔCt method (Livak and Schmittgen, 2001). For dot blots, 1 µg and 2 µg of total RNA were spotted onto a Hybond-N+ (Amersham Bioscience), and dsRNAs were immunodetected with the anti-dsRNA J2 antibody (Jena Bioscience) as previously described (Baudry *et al*., 2026). Primers and probes used are listed in Table S1.

### Immunoblot analysis

Proteins were extracted in a ratio of 450 µL of 2x Laemmli containing 15 mg/mL DTT per 100mg of tissue and separated by 10% SDS-PAGE. The Flag tag was immunodetected with anti-Flag F3165 (Sigma) as previously described (Baudry *et al*., 2026).

## AUTHOR CONTRIBUTIONS

Sarah Gartner (Data curation, Formal analysis, Investigation, Visualization, Writing); Clara Bougearel (Investigation); Gadi Schuster (Conceptualization, Funding acquisition, Project administration, Writing); Benoît Castandet (Conceptualization, Funding acquisition, Project administration, Writing); David B. Stern (Conceptualization, Funding acquisition, Project administration, Writing); Kevin Baudry (Conceptualization, Data curation, Formal analysis, Investigation, Project administration, Visualization, Writing). All authors participated in Writing – review & editing.

## SUPPORTING INFORMATION

**Figure S1.** Localization of RNase J signature motifs and motif II conservation.

**Figure S2.** Transgene copy number is not correlated with the *rnj-2* zygosity background.

**Figure S3.** Characterization of the progeny of a m2ΔGT-1 chlorotic plant.

**Figure S4.** Time course of RNJ accumulation in m2ΔGT-1.

**Figure S5.** RNJ abundance in plants dissected at different stage of chlorosis in m2ΔGT-1.

**Figure S6**. Chlorosis development and transgene copy number in WTox lines.

**Figure S7.** Molecular comparison of green seedlings and chlorotic adult WTox plants.

**Table S1.** Primers and probes used in this article

## REFERENCES

Bao, L., Hu, J., Zhan, B., et al. (2023) Structural insights into RNase J that plays an essential role in Mycobacterium tuberculosis RNA metabolism. Nature Communications 2023 14:1, 14, 1–14. Available at: https://www.nature.com/articles/s41467-023-38045-z [Accessed October 2, 2025].

Baudry, K., Delannoy, E. and Colas des Francs-Small, C. (2022) Analysis of the Plant Mitochondrial Transcriptome. Methods in Molecular Biology, 2363, 235–262. Available at: https://link.springer.com/protocol/10.1007/978-1-0716-1653-6_17 [Accessed October 2, 2025].

Baudry, K., Skiada, S., Burstein, C., et al. (2026) The GT1 domain of RNase J ensures RNA quality control through dsRNA binding in Arabidopsis plastids. Nucleic Acids Res., 54, 13–14. Available at: 10.1093/nar/gkag033 [Accessed March 22, 2026].

Bollenbach, T.J., Lange, H., Gutierrez, R., Erhardt, M., Stern, D.B. and Gagliardi, D. (2005) RNR1, a 3’-5’ exoribonuclease belonging to the RNR superfamily, catalyzes 3’ maturation of chloroplast ribosomal RNAs in Arabidopsis thaliana. Nucleic Acids Res., 33, 2751–2763. Available at: http://nar.oupjournals.org/cgi/reprint/33/8/2751?ijkey=E6tjwhBeaJDrp7x&keytype=ref.

Castandet, B., Germain, A., Hotto, A.M. and Stern, D.B. (2019) Systematic sequencing of chloroplast transcript termini from Arabidopsis thaliana reveals >200 transcription initiation sites and the extensive imprints of RNA-binding proteins and secondary structures. Nucl. Acids Res., 47, 11889–11905.

Castandet, B., Hotto, A.M., Fei, Z. and Stern, D.B. (2013) Strand-specific RNA sequencing uncovers chloroplast ribonuclease functions. FEBS Lett, 587, 3096–3101. Available at: http://www.ncbi.nlm.nih.gov/pubmed/23954624.

Chen, H., Zou, W. and Zhao, J. (2015) Ribonuclease J is required for chloroplast and embryo development in Arabidopsis. J. Exp. Bot., 66, 2079–2091. Available at: http://jxb.oxfordjournals.org/content/early/2015/02/20/jxb.erv010.abstract.

Chen, J., Zhu, H., Huang, J. and Huang, W. (2020) A new method for functional analysis of plastid EMBRYO-DEFECTIVE PPR genes by efficiently constructing cosuppression lines in Arabidopsis. Plant Methods, 16, 154-. Available at: https://link.springer.com/article/10.1186/s13007-020-00696-0 [Accessed August 30, 2026].

Chen, L., Chen, J.Y., Zhang, X., et al. (2017) R-ChIP Using Inactive RNase H Reveals Dynamic Coupling of R-loops with Transcriptional Pausing at Gene Promoters. Mol. Cell, 68, 745. Available at: https://pmc.ncbi.nlm.nih.gov/articles/PMC5957070/ [Accessed August 24, 2026].

Cheng, Z.F. and Deutscher, M.P. (2002) Purification and characterization of the Escherichia coli exoribonuclease RNase R. Comparison with RNase II. J. Biol. Chem., 277, 21624–21629. Available at: http://www.ncbi.nlm.nih.gov/entrez/query.fcgi?cmd=Retrieve&db=PubMed&dopt=Citation&list_uids=11948193.

Clough, S.J. and Bent, A.F. (1998) Floral dip: a simplified method for Agrobacterium-mediated transformation of Arabidopsis thaliana. Plant J., 16, 735–743. Available at: http://www.ncbi.nlm.nih.gov/entrez/query.fcgi?cmd=Retrieve&db=PubMed&dopt=Citation&list_uids=10069079.

Daou-Chabo, R. and Condon, C. (2009) RNase J1 endonuclease activity as a probe of RNA secondary structure. RNA, 15, 1417–1425. Available at: http://www.ncbi.nlm.nih.gov/entrez/query.fcgi?cmd=Retrieve&db=PubMed&dopt=Citation&list_uids=19458035.

Dorléans, A., Li de la Sierra-Gallay, Inés, Piton, J., et al. (2011) Molecular basis for the recognition and cleavage of RNA by the bifunctional 5’-3’ exo/endoribonuclease RNase J. Structure, 19, 1252–1261.

Ganusova, E.E., Reagan, B.C., Fernandez, J.C., et al. (2020) Chloroplast-to-nucleus retrograde signalling controls intercellular trafficking via plasmodesmata formation. Philosophical Transactions of the Royal Society B: Biological Sciences, 375. Available at: 10.1098/rstb.2019.0408 [Accessed August 27, 2026].

Gatewood, M.L., Bralley, P., Weil, M.R. and Jones, G.H. (2012) RNA-Seq and RNA Immunoprecipitation Analyses of the Transcriptome of Streptomyces coelicolor Identify Substrates for rnase III. J. Bacteriol., 194, 2228–2237.

Georg, J., Honsel, A., Voss, B., Rennenberg, H. and Hess, W.R. (2010) A long antisense RNA in plant chloroplasts. New Phytol. Available at: http://www.ncbi.nlm.nih.gov/entrez/query.fcgi?cmd=Retrieve&db=PubMed&dopt=Citation&list_uids=20202127.

Germain, A., Herlich, S., Larom, S., Kim, S.H., Schuster, G. and Stern, D.B. (2011) Mutational analysis of Arabidopsis chloroplast polynucleotide phosphorylase reveals roles for both RNase PH core domains in polyadenylation, RNA 3’-end maturation and intron degradation. Plant J., 67, 381–394. Available at: http://www.ncbi.nlm.nih.gov/pubmed/21466602.

Germain, A., Kim, S.H., Gutierrez, R. and Stern, D.B. (2012) Ribonuclease II preserves chloroplast RNA homeostasis by increasing mRNA decay rates, and cooperates with polynucleotide phosphorylase in 3’ end maturation. Plant J., 72, 960–971. Available at: http://www.ncbi.nlm.nih.gov/pubmed/23061883.

Halpert, M., Liveanu, V., Glaser, F. and Schuster, G. (2019) The Arabidopsis chloroplast RNase J displays both exo- and robust endonucleolytic activities. Plant Mol Biol, 99, 17–29.

Hong, Z.H., Zhu, L., Gao, L.L., Zhu, Z., Su, T., Krall, L., Wu, X.N., Bock, R. and Wu, G.Z. (2025) Chloroplast precursor protein preClpD overaccumulation triggers multilevel reprogramming of gene expression and a heat shock-like response. Nature Communications 2025 16:1, 16, 3777-. Available at: https://www.nature.com/articles/s41467-025-59043-3 [Accessed September 27, 2026].

Hotto, A.M., Huston, Z.E. and Stern, D.B. (2010) Overexpression of a natural chloroplast-encoded antisense RNA in tobacco destabilizes 5S rRNA and retards plant growth. BMC Plant Biol., 10, 213. Available at: http://www.ncbi.nlm.nih.gov/entrez/query.fcgi?cmd=Retrieve&db=PubMed&dopt=Citation&list_uids=20920268.

Hotto, A.M., Schmitz, R.J., Fei, Z., Ecker, J.R. and Stern, D.B. (2011) Unexpected diversity of chloroplast noncoding RNAs as revealed by deep sequencing of the Arabidopsis transcriptome. G3, 1, 559–570. Available at: http://www.g3journal.org/content/1/7/559.abstract.

Jan, M., Liu, Z., Rochaix, J.D. and Sun, X. (2022) Retrograde and anterograde signaling in the crosstalk between chloroplast and nucleus. Front. Plant Sci., 13, 980237. Available at: [Accessed August 31, 2026].

Jeran, N., Mercier, M., Pesaresi, P. and Tadini, L. (2025) Proteostasis and protein quality control in chloroplasts: mechanisms and novel insights related to protein mislocalization. J. Exp. Bot., 76, 4294–4312. Available at: 10.1093/jxb/eraf182 [Accessed September 27, 2026].

Kiselev, K. V., Aleynova, O.A., Ogneva, Z. V., Suprun, A.R. and Dubrovina, A.S. (2021) 35S promoter-driven transgenes are variably expressed in different organs of Arabidopsis thaliana and in response to abiotic stress. Mol. Biol. Rep., 48, 2235–2241. Available at: https://link.springer.com/article/10.1007/s11033-021-06235-x [Accessed August 30, 2026].

Lanet, E., Delannoy, E., Sormani, R., Floris, M., Brodersen, P., Crété, P., Voinnet, O. and Robaglia, C. (2009) Biochemical Evidence for Translational Repression by Arabidopsis MicroRNAs. Plant Cell, 21, 1762. Available at: https://pmc.ncbi.nlm.nih.gov/articles/PMC2714937/ [Accessed August 30, 2026].

Levy, S., Portnoy, V., Admon, J. and Schuster, G. (2011) Distinct activities of several RNase J proteins in methanogenic archaea. RNA Biol., 8, 1073–1083.

Lioliou, E., Sharma, C.M., Caldelari, I., Helfer, A.C., Fechter, P., Vandenesch, F., Vogel, J. and Romby, P. (2012) Global regulatory functions of the Staphylococcus aureus endoribonuclease III in gene expression. PLoS Genet., 8.

Liponska, A., Jamalli, A., Kuras, R., et al. (2018) Tracking the elusive 5 ′ exonuclease activity of Chlamydomonas reinhardtii RNase J. Plant Mol. Biol., 96, 641–653.

Livak, K.J. and Schmittgen, T.D. (2001) Analysis of Relative Gene Expression Data Using Real-Time Quantitative PCR and the 2−ΔΔCT Method. Methods, 25, 402–408. Available at: [Accessed September 1, 2026].

Luro, S., Germain, A., Sharwood, R.E. and Stern, D.B. (2013) RNase J participates in a pentatricopeptide repeat protein-mediated 5’ end maturation of chloroplast mRNAs. Nucleic Acids Res, 41, 9141–9151. Available at: http://www.ncbi.nlm.nih.gov/pubmed/23921629.

Mathy, N., Benard, L., Pellegrini, O., Daou, R., Wen, T. and Condon, C. (2007) 5’-to-3’ exoribonuclease activity in bacteria: role of RNase J1 in rRNA maturation and 5’ stability of mRNA. Cell, 129, 681–692. Available at: http://www.ncbi.nlm.nih.gov/entrez/query.fcgi?cmd=Retrieve&db=PubMed&dopt=Citation&list_uids=17512403.

Mathy, N., Hebert, A., Mervelet, P., Benard, L., Dorleans, A., Li de la Sierra-Gallay, I., Noirot, P., Putzer, H. and Condon, C. (2010) Bacillus subtilis ribonucleases J1 and J2 form a complex with altered enzyme behaviour. Mol. Microbiol., 75, 489–498. Available at: http://www.ncbi.nlm.nih.gov/pubmed/20025672.

Newman, J.A., Hewitt, L., Rodrigues, C., Solovyova, A., Harwood, C.R. and Lewis, R.J. (2011) Unusual, dual endo- and exonuclease activity in the degradosome explained by crystal structure analysis of RNase J1. Structure, 19, 1241–1251. Available at: http://www.ncbi.nlm.nih.gov/pubmed/21893285.

Pei, X.-Y.Y., Bralley, P., Jones, G.H. and Luisi, B.F. (2015) Linkage of catalysis and 5’ end recognition in ribonuclease RNase J. Nucleic Acids Res, 43, 8066–8076.

Perrin, R., Meyer, E.H., Zaepfel, M., Kim, Y.J., Mache, R., Grienenberger, J.M., Gualberto, J.M. and Gagliardi, D. (2004) Two exoribonucleases act sequentially to process mature 3’-ends of atp9 mRNAs in Arabidopsis mitochondria. J. Biol. Chem., 279, 25440–25446. Available at: http://www.ncbi.nlm.nih.gov/entrez/query.fcgi?cmd=Retrieve&db=PubMed&dopt=Citation&list_uids=15037609.

Pfalz, J., Bayraktar, O.A., Prikryl, J. and Barkan, A. (2009) Site-specific binding of a PPR protein defines and stabilizes 5’ and 3’ mRNA termini in chloroplasts. EMBO J., 28, 2042–2052. Available at: http://www.ncbi.nlm.nih.gov/entrez/query.fcgi?cmd=Retrieve&db=PubMed&dopt=Citation&list_uids=19424177.

Qi, Y., Armbruster, U., Schmitz-Linneweber, C., Delannoy, E., Longevialle, A.F. de, Ruhle, T., Small, I., Jahns, P. and Leister, D. (2011) Arabidopsis CSP41 proteins form multimeric complexes that bind and stabilize distinct plastid transcripts. J. Exp. Bot. Available at: http://www.ncbi.nlm.nih.gov/pubmed/22090436.

Raj, R., Nadig, S., Patel, T. and Gopal, B. (2020) Structural and biochemical characteristics of two Staphylococcus epidermidis RNase J paralogs RNase J1 and RNase J2. Journal of Biological Chemistry, 295, 16863–16876. Available at: https://www.jbc.org/action/showFullText?pii=S0021925817504986 [Accessed October 2, 2025].

Rovira, A.G. and Smith, A.G. (2019) PPR proteins – orchestrators of organelle RNA metabolism. Physiol. Plant., 166, 451–459. Available at: http://www.ncbi.nlm.nih.gov/pubmed/30809817 [Accessed August 13, 2019].

Schmid, L.M., Manavski, N., Chi, W. and Meurer, J. (2024) Chloroplast Ribosome Biogenesis Factors. Plant Cell Physiol., 65, 516–536. Available at: 10.1093/pcp/pcad082 [Accessed August 26, 2026].

Sharwood, R.E., Halpert, M., Luro, S., Schuster, G. and Stern, D.B. (2011) Chloroplast RNase J compensates for inefficient transcription termination by removal of antisense RNA. RNA, 17, 2165–2176.

Small, I.D., Schallenberg-Rüdinger, M., Takenaka, M., Mireau, H. and Ostersetzer-Biran, O. (2020) Plant organellar RNA editing: what 30 years of research has revealed. The Plant Journal, 101, 1040–1056. Available at: https://onlinelibrary.wiley.com/doi/full/10.1111/tpj.14578 [Accessed April 28, 2026].

Stoppel, R. and Meurer, J. (2012) Complex RNA metabolism in the chloroplast: an update on the psbB operon. Planta. Available at: http://www.ncbi.nlm.nih.gov/pubmed/23065055.

Tadini, L., Jeran, N., Domingo, G., et al. (2023) Perturbation of protein homeostasis brings plastids at the crossroad between repair and dismantling. PLoS Genet., 19, e1010344. Available at: https://journals.plos.org/plosgenetics/article?id=10.1371/journal.pgen.1010344 [Accessed September 27, 2026].

Tejada-Arranz, A., Lulla, A., Bouilloux-Lafont, M., et al. (2023) Acetylation regulates the oligomerization state and activity of RNase J, the Helicobacter pylori major ribonuclease. Nature Communications 2023 14:1, 14, 1–15. Available at: https://www.nature.com/articles/s41467-023-43825-8 [Accessed October 2, 2025].

Walter, M., Kilian, J. and Kudla, J. (2002) PNPase activity determines the efficiency of mRNA 3’-end processing, the degradation of tRNA and the extent of polyadenylation in chloroplasts. EMBO J., 21, 6905–6914. Available at: http://www.ncbi.nlm.nih.gov/entrez/query.fcgi?cmd=Retrieve&db=PubMed&dopt=Citation&list_uids=12486011.

Walter, M., Piepenburg, K., Schöttler, M.A., et al. (2010) Knockout of the plastid RNase E leads to defective RNA processing and chloroplast ribosome deficiency. Plant J., 64, 851–863. Available at: 10.1111/j.1365-313X.2010.04377.x.

Wang, X., Wang, J., Li, S., Lu, C. and Sui, N. (2022) An overview of RNA splicing and functioning of splicing factors in land plant chloroplasts. RNA Biol., 19, 897–907. Available at: https://www.tandfonline.com/doi/abs/10.1080/15476286.2022.2096801 [Accessed April 26, 2026].

Yu, F., Fu, A., Aluru, M., et al. (2007) Variegation mutants and mechanisms of chloroplast biogenesis. Plant Cell Environ., 30, 350–365. Available at: https://onlinelibrary.wiley.com/doi/full/10.1111/j.1365-3040.2006.01630.x [Accessed August 25, 2026].

Zghidi-Abouzid, O., Merendino, L., Buhr, F., Malik Ghulam, M. and Lerbs-Mache, S. (2011) Characterization of plastid psbT sense and antisense RNAs. Nucleic Acids Res., 39, 5379– 5387.

Zhao, Y., Lu, M., Zhang, H., et al. (2015) Structural insights into catalysis and dimerization enhanced exonuclease activity of RNase J. Nucleic Acids Res., 43, 5550–5559. Available at: https://academic.oup.com/nar/article-lookup/doi/10.1093/nar/gkv444 [Accessed April 4, 2019].

Zhelyazkova, P., Sharma, C.M., Forstner, K.U., Liere, K., Vogel, J. and Börner, T. (2012) The primary transcriptome of barley chloroplasts: numerous noncoding RNAs and the dominating role of the plastid-encoded RNA polymerase. Plant Cell, 24, 123–136. Available at: http://www.ncbi.nlm.nih.gov/pubmed/22267485.

Zheng, X., Feng, N., Li, D., Dong, X. and Li, J. (2017) New molecular insights into an archaeal RNase J reveal a conserved processive exoribonucleolysis mechanism of the RNase J family. Mol. Microbiol., 106, 351–366.

