## Supplemental_Figures-Table for "Overexpression of Active or Inactive RNase J Disrupts Chloroplast RNA Quality Control in Arabidopsis"

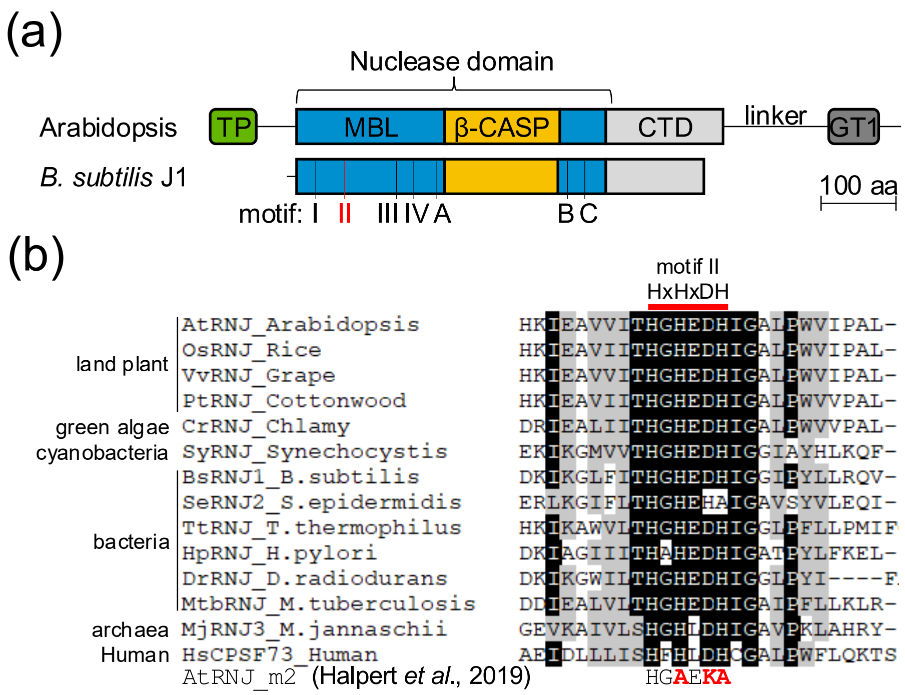


**Figure S1.** Localization of RNase J signature motifs and motif II conservation.
(a) RNJ linear structures from *Arabidopsis thaliana* and *Bacillus subtilis.* Domains drawn to scale are the TP, chloroplast target peptide; MBL, Metallo-β-lactamase domain; β-CASP, β-CASP domain; CTD, C-terminal domain; GT1, GT1 domain. At the bottom, MBL signature motifs are indicated (I-IV, A-C).
(b) Protein alignment of 14 RNJ orthologs showing motif II (aa 175-180 in Arabidopsis). Below, residues in red show the mutated version of AtRNJ motif II used in Halpert *et al.*, 2019 and in this article. Uniprot IDs: AtRNJ: Q84W56, OsRNJ:A2X5S9, VvRNJ: A0ABY9DJN0, PtRNJ: B9I3P4, CrRNJ: B2YFW5, SyRNJ: P54123, BsRNJ1: Q45493, SeRNJ2: Q8CST0, TtRNJ: Q72JJ7, HpRNJ: P56185, DrRNJ: H9CZL7, MtbRNJ: P9WGZ9, MjRNJ3: Q58271, HsCPSF73: Q9UKF6.

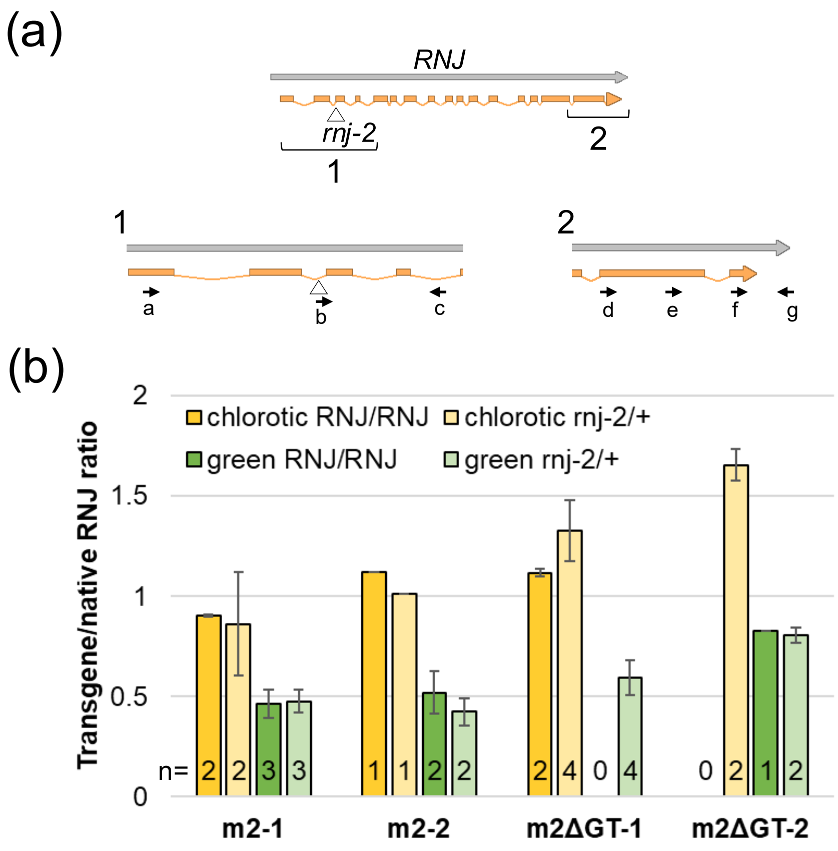


**Figure S2.** Transgene copy number is not correlated with the *rnj-2* zygosity background.
(a) Gene model of *RNJ* gene, which is disrupted by a T-DNA in *rnj-2* as indicated by the triangle. Regions indicated by brackets are magnified below to show primer binding sites. Region 1 shows primers used to genotype the *rnj-2* T-DNA insertion, with the wild-type allele being amplified with primers a and c, and the T-DNA insertion with primers b and c. Region 2 shows primers used to measure *RNJ* expression or copy number, with f and g being used to measure the native *RNJ* gene. Transgenes were measured with a primer in the Flag tag and primer e (full-length constructs) or d (m2ΔGT).
(b) Same qPCR data as shown in Figure 3, but distinguishing between different genetic backgrounds. Chlorotic plants are shown in yellow, and green plants in green; WT (*RNJ/RNJ*) and heterozygous (*rnj-2/+*) backgrounds are depicted by darker and lighter hues, respectively. Number of plants in each bin (n) is shown at the bottom of each bar.

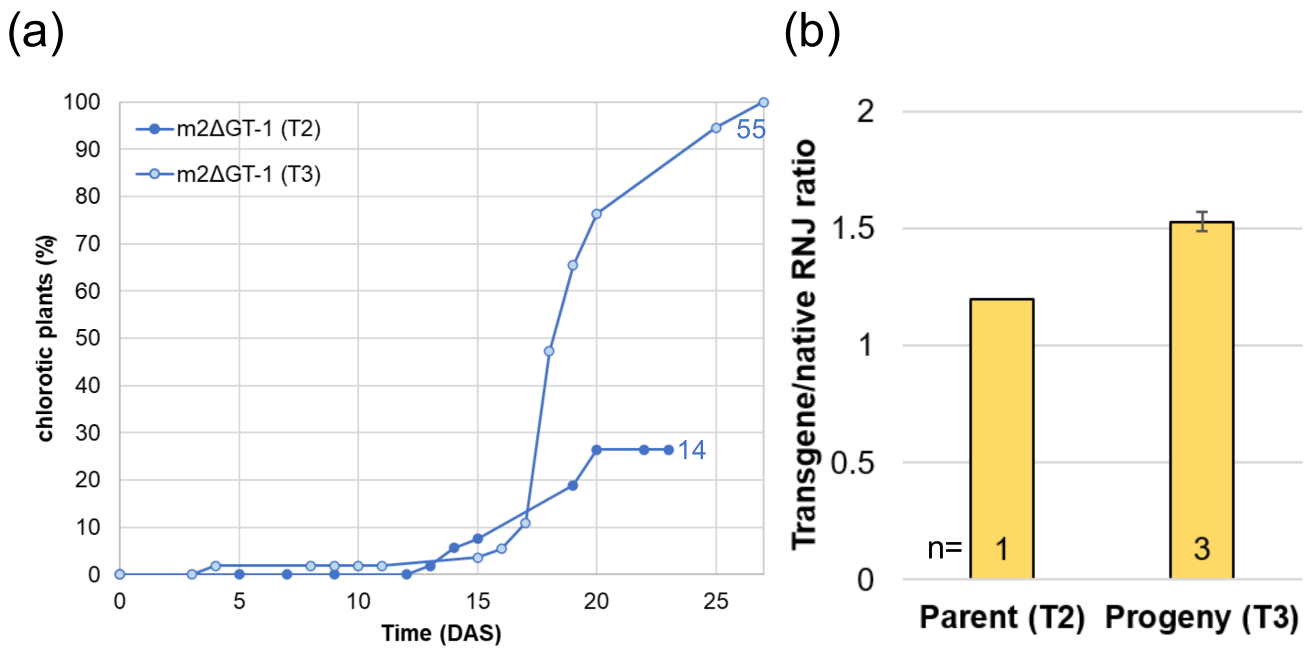


**Figure S3.** Characterization of the progeny of a m2ΔGT-1 chlorotic plant.
(a) Percentage of chlorotic plants in the T3 generation of the m2ΔGT-1 line at different time points (DAS: days after sowing). Curve for the T2 generation shown in Figure 2 is added for comparison.
(b) Transgene copy number evaluation by qPCR in a chlorotic T2 plant (Parent) and in 3 randomly chosen T3 plants among its progeny (Progeny). Number of plants analyzed for each bin (n) is shown at the bottom of each bar.

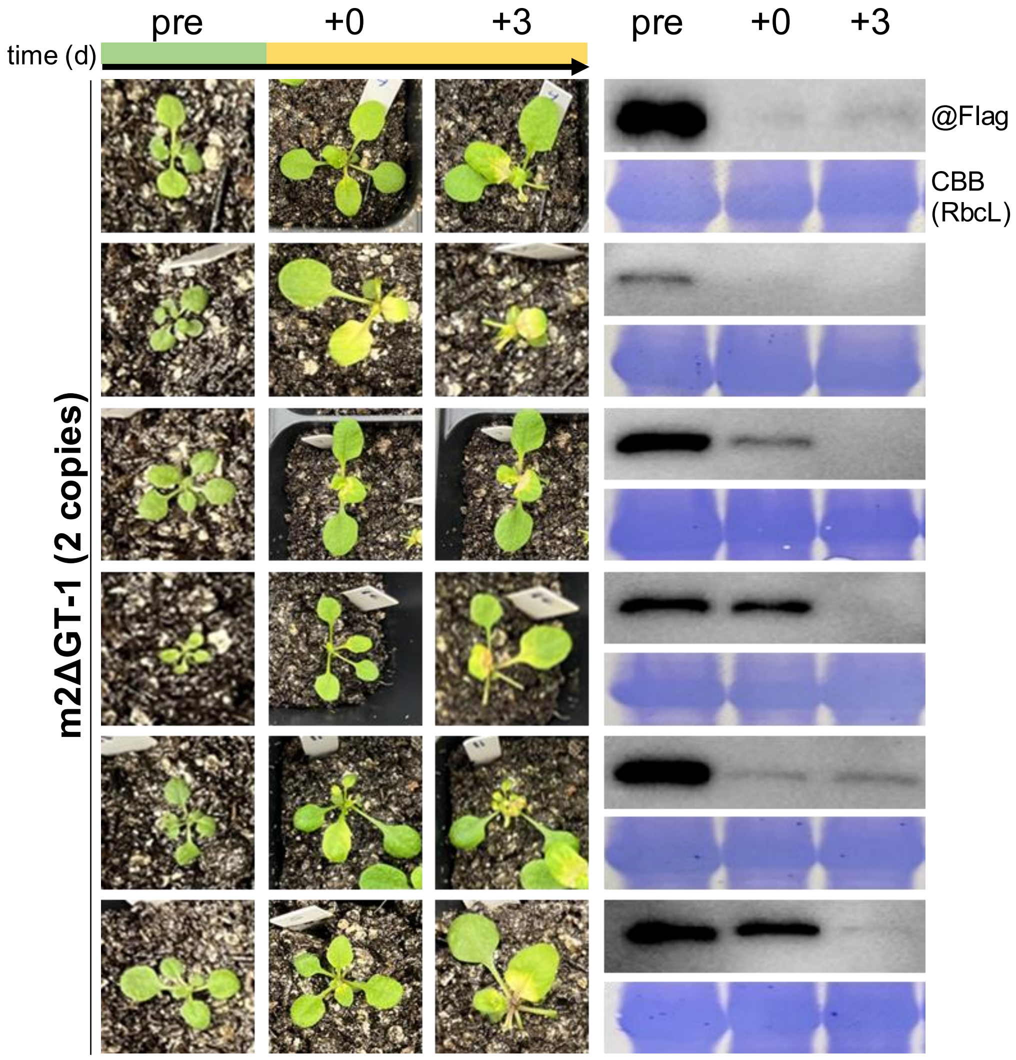


**Figure S4.** Time course of RNJ accumulation in m2ΔGT-1.
Left: Time course of plants containing two transgene copies before chlorosis appears (pre), on the first day the chlorosis was noted (+0), and three days later. For each series, the same plant is followed over time and leaf tissue was collected at each time point. Right: Protein analysis of the associated sample at each time point as indicated at the top. Top, immunoblot with anti-Flag antibody showing presence/absence of the m2ΔGT protein. Lower, Coomassie-stained gel cropped to show the region of the Rubisco large subunit.

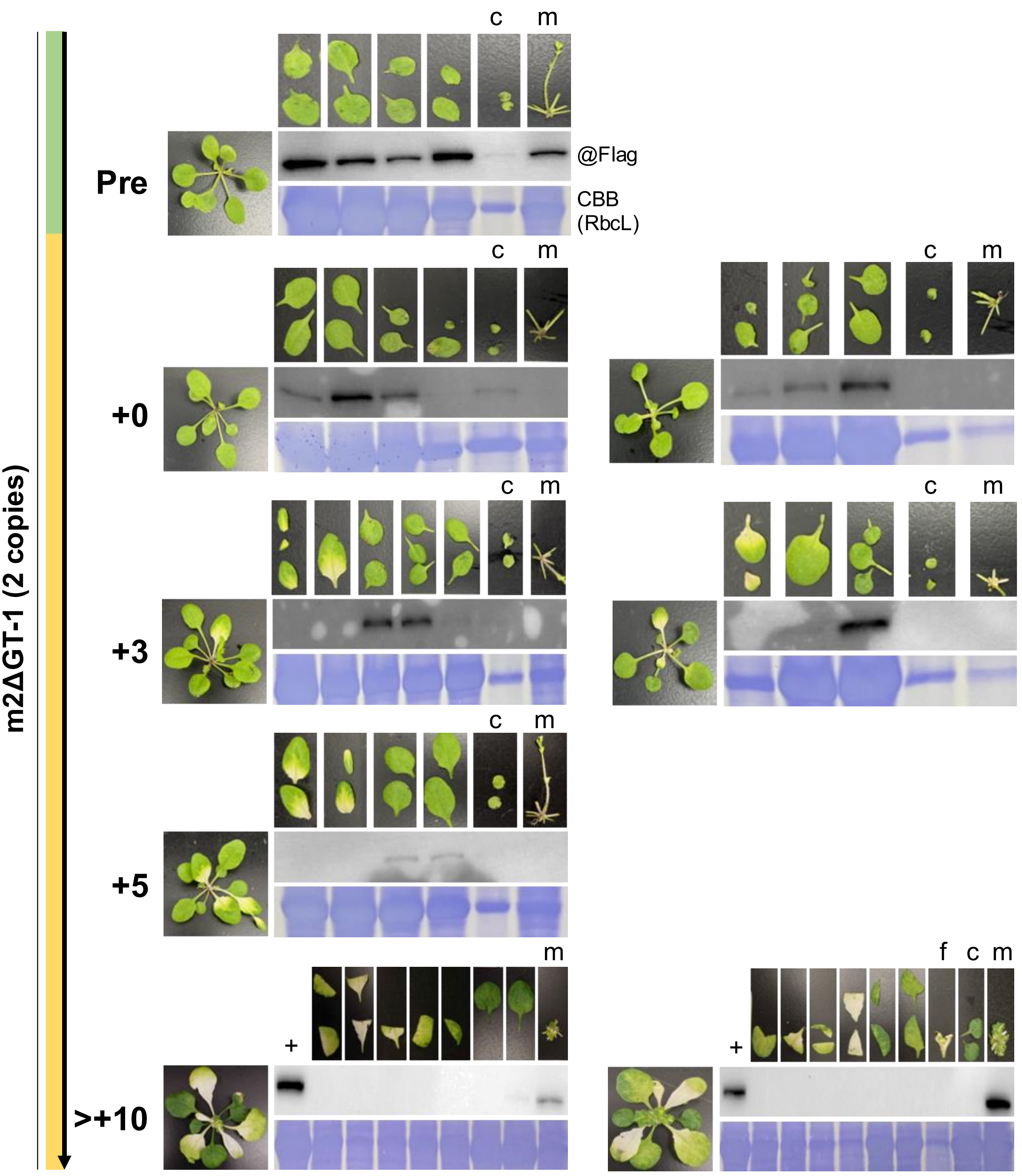


**Figure S5.** RNJ abundance in plants dissected at different stage of chlorosis in m2ΔGT-1.
RNJ accumulation in dissected plants containing two copies of the transgene before chlorosis appears (pre), on the first day the chlorosis was noted (+0), 3 and 5 days, and more than 10 days after. For pre and +5, one plant is shown, whereas two are shown for +0, +3 and >+10 time points. For each series, the full plant before being dissected is shown on the left of the blot. The collected tissue is shown above each lane, “c” indicates cotyledon, “f” indicates white flower bud, “+” indicates a positive control coming from a green leaf from a green “1 copy” plant, “m” indicates the middle and remaining part of the rosette after cutting out all leaves.

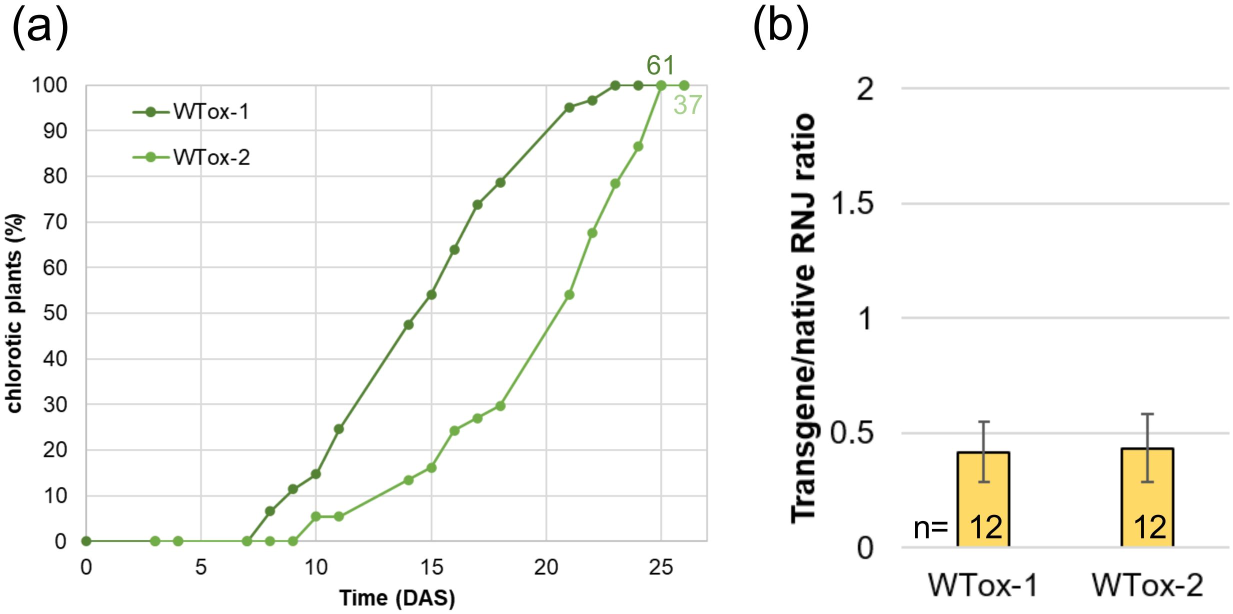


**Figure S6**. Chlorosis development and transgene copy number in WTox lines.
(a) Percentage of chlorotic plants of the indicated WTox lines at different time points (DAS: days after sowing). Numbers at the end of each curve represent the total number of chlorotic plants observed.
(b) Transgene copy number evaluation by qPCR in WTox lines. The number of plants analyzed for each bin (n) is shown at the bottom of each bar.

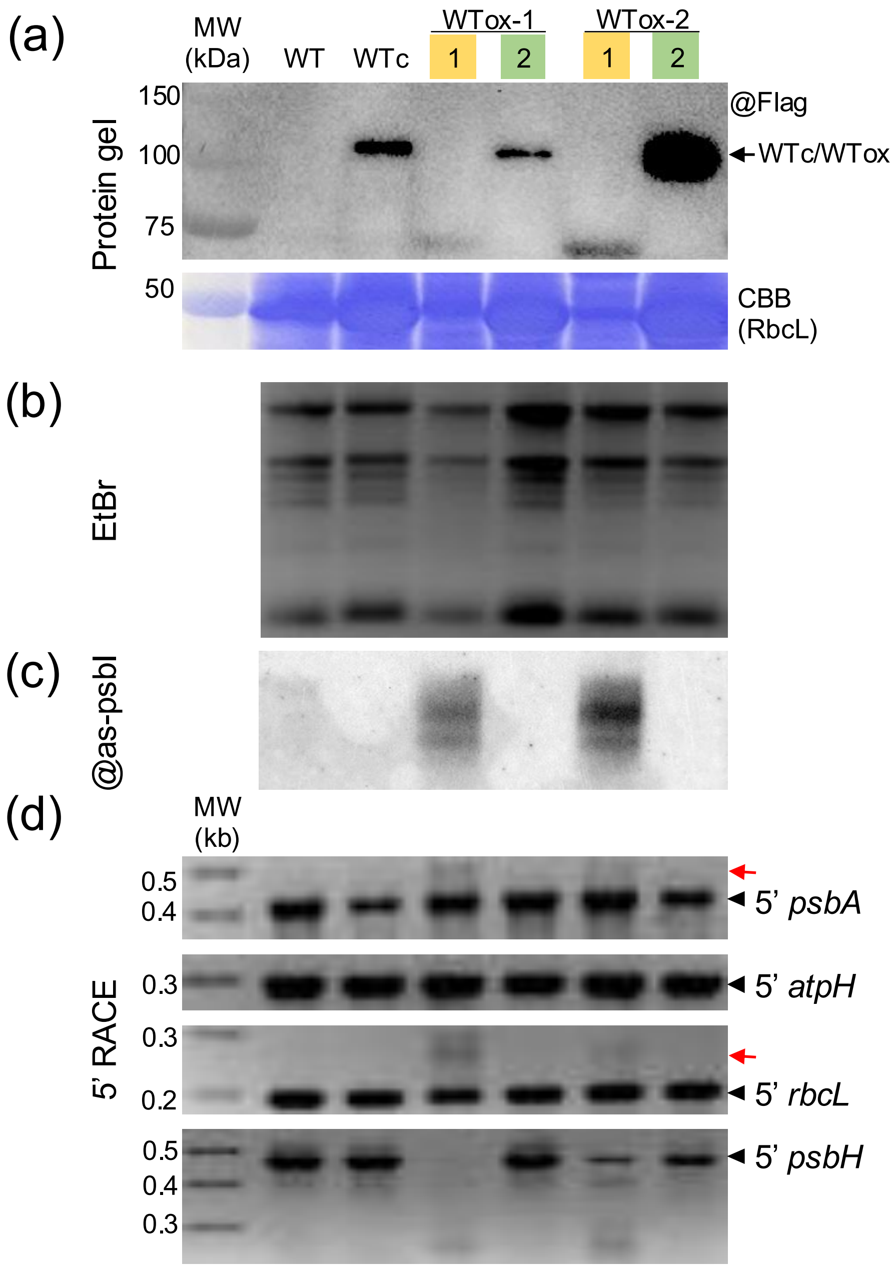


**Figure S7.** Molecular comparison of green seedlings and chlorotic adult WTox plants.
Green and yellow boxes indicate 2 week old still green seedling and 4 week old adult chlorotic plants, respectively. Sizes in kb are shown to the left of gels.
(a) Protein analysis from the indicated lines.
(b) Ethidium bromide-stained gel of 4 µg total RNA from the same lines as in panel (a).
(c) RNA gel blot detection of *as-psbI* from the same samples as in panel (b).
(d) 5' RACE analysis from the same samples as in panel (b).

**Table S1.** Primers and Probes used in this article

| Name | Sequence | Comments |
|---|---|---|
| **Genotyping** | | |
| rnj2_LP | CCCTCTCGCTTTGTCCCTAT | rnj-2 genotyping |
| rnj2_RP | CGCAAGCTGAGGTACATACA | rnj-2 genotyping |
| LB1 | GCCTTTTCAGAAATGGATAAATAGCCTTGCTTCC | rnj-2 genotyping |
| AtRNJ_ex16_F | ACTGCGGCATCAACATACAC | transgene genotyping |
| FLAGtag_R | TGTCATCGTCATCCTTGTAGTC | transgene genotyping |
| **(RT-)qPCR** | | |
| AtActin1_F | TCTTGATCTTGCTGGTCGTG | ACT1 (At2g37620) gene |
| AtActin1_R | GAGCTGGTTTTGGCTGTCTC | ACT1 (At2g37620) gene |
| AtRNJ_ex17_qPCR_F | GAGAAGCAAGACGAGTTGGC | native RNJ (At5g63420) gene |
| AtRNJ_3UTR_qPCR_R | GGGATTTGGATTCTGCGTTGA | native RNJ (At5g63420) gene |
| AtRNJ_GT1_qPCR_F | GAGGAGAGCTGCACAGTAGA | m2, WTox, WTc transgene |
| FLAGtag_qPCR_R | AGCCTTGTCATCGTCATCCT | m2, WTox, WTc transgene |
| AtRNJ_ex16_F | ACTGCGGCATCAACATACAC | m2ΔGT transgene |
| FLAGtag_R | TGTCATCGTCATCCTTGTAGTC | m2ΔGT transgene |
| LHCB1.2_qPCR_F | GGACTTGCTTTACCCCGGTG | LHCB1.2 (At1g29910) gene |
| LHCB1.2_qPCR_R | TCGGTAGCAAGACCCAATGG | LHCB1.2 (At1g29910) gene |
| GLK1_qPCR_F | CGGAGCGTCAGAGTTTCTTG | GLK1 (At2g20570) gene |
| GLK1_qPCR_R | CCTTTCCCCGAACTACCCTT | GLK1 (At2g20570) gene |
| AtUBC21_qPCR_F | CTGCGACTCAGGGAATCTTCTA | UBC21 (At5g25760) gene |
| AtUBC21_qPCR_R | TTGTGCCATTGAATTGAACCC | UBC21 (At5g25760) gene |
| **5'RACE** | | |
| 5'RACEAdapter | AUAUGCGCGAAUUCCUGUAGAACGAACACUAGAAGAAA | RNA adapter ligated |
| ASP | AATTCCTGTAGAACGAACACTAG | Adaptor Specific Primer |
| 5'RACE_AtPsbA | AGCCATTCATCAACGGATGC | amplify psbA 5' |
| 5'RACE_AtAtpH | GCGCTAATGCTACAACCAGGCC | amplify atpH 5' |
| 5'RACE_AtRbcL | TCCTTGGTTTCATATTCAGGA | amplify rbcL 5' |
| 5'RACE_AtPsbH | GGCTAAAATTGTTAGACCTGCAT | amplify psbH 5' |
| **Northern Blot** | | |
| BioProbe_AtPsbI-as | bio-CTCTCAAACTTTTTGTATACACTGTAGTTATATTCTTTGT | biotinylated oligonucleotide to detect antisense-psbI |
